# Genetic engineering for *in vivo* optical interrogation of neuronal responses to cell type-specific silencing

**DOI:** 10.1101/2021.01.13.426508

**Authors:** Firat Terzi, Johannes Knabbe, Sidney B. Cambridge

**Affiliations:** Department of Functional Neuroanatomy, Institute of Cell Biology and Anatomy, Heidelberg University, Im Neuenheimer Feld 307, 69120 Heidelberg, Germany

**Keywords:** Conditional transgene induction, Cre/lox system, Tet system, in vivo two-photon microscopy, Kir2.1, neuronal silencing, homeostasis

## Abstract

Genetic engineering of quintuple transgenic brain tissue was used to establish a low background, Cre-dependent version of the inducible Tet-On system for fast, cell type-specific transgene expression *in vivo*. Co-expression of a constitutive, Cre-dependent fluorescent marker selectively allowed single cell analyses *before* and *after* inducible, tet-dependent transgene expression. Here, we used this method for acute, high-resolution manipulation of neuronal activity in the living brain. Single induction of the potassium channel Kir2.1 produced cell type-specific silencing within hours that lasted for at least three days. Longitudinal *in vivo* imaging of spontaneous calcium transients and neuronal morphology demonstrated that prolonged silencing did not alter spine densities or synaptic input strength. Furthermore, selective induction of Kir2.1 in parvalbumin interneurons increased the activity of surrounding neurons in a distance-dependent manner. This high-resolution, inducible interference and interval imaging of individual cells (high I^5^, ‘HighFive’) method thus allows visualizing temporally precise, genetic perturbations of defined cells.

## Introduction

At the level of individual cells, the phenotypic consequences of genetic perturbations are difficult to assess in whole organs. Potentially subtle changes in cellular phenotypes after overexpression or knock-down/out of genes are sometimes hard to detect on a tissue level, let alone in specific cell types. Particularly in the brain with hundreds of different types of neurons (Saunders et al., 2018), it is not surprising that the contributions of many genes to neuronal function remain unclear. Of course, neuronal plasticity and homeostatic mechanisms only aggravate this problem as perturbations to the brain are often compensated for on various time-scales (Pozo and Goda, 2010; Turrigiano, 2011). We thus need to be able to manipulate genes rapidly and identify their acute phenotypic effects in clearly defined cell populations. Additionally, it would be highly advantageous to be able to characterize secondary or compensatory effects of cells neighboring the manipulated cells. However, despite decades of progress in the field of conditional transgene expression, rapid, cell type-specific transgene induction remains difficult to achieve with currently available methods. The powerful Cre/lox system is widely used for expression of transgenes in specific cell-types (Yarmolinsky and Hoess, 2015). Cellular specificity is achieved through well-characterized endogenous or exogenous promoters. The conditional Tamoxifen-inducible Cre/lox version (Feil et al., 1996) adds temporal control although the time resolution is poor as transgene expression is typically induced by injections of Tamoxifen over several days. Alternatively, the tetracycline (Tet) system is a two-component system (Mansuy and Bujard, 2000) (transactivator + tet-dependent transgene) that is also commonly used for conditional transgene expression but there are much fewer cell type-specific promoter constructs available for brain research. Transgene expression with the Tet-Off system is considerably slow, because induction occurs once doxycycline is physiologically cleared from the animal which takes at least a few days (Kistner et al., 1996). Conversely, the faster Tet-On system was shown to express transgenes within hours after administration (Hasan et al., 2001), but there tend to be issues with elevated background expression and lower inducibility (Zhu et al., 2007). Thus, both, the Cre/lox and the Tet systems have strengths and weaknesses. Moreover, there are currently no methods existing that allow prior identification and visualization of those cells in which the respective transgene will be induced. Of course, this would be very desirable for live analyses and optical interrogation of the acute effects of transgene expression on cells and tissues. We achieved this by rendering the fast, inducible Tet-On system (Dogbevia et al., 2015; Gossen et al., 1995) dependent on the cell type-specific, but low-temporal resolution Cre recombinase system (Pluck, 1996). Our approach combines the strengths of both transgene expression methods by providing fast, tet-dependent expression which is restricted by the cellular specificity of the Cre/lox system. In addition, Cre-mediated constitutive co-expression of a fluorescent reporter highlights cells that are primed for induction of a Cre- and tet-dependent transgene. By directly expressing transgenes that cell-autonomously change the activity of neurons, we used longitudinal *in vivo* two-photon microscopy to demonstrate that one can use this approach as an attractive complement to the DREADD system (Armbruster et al., 2007) or optogenetics (Emiliani et al., 2015), especially for long-term manipulation in the range of a few days.

## Results

To develop an inducible approach for fast and cell type-specific expression in the living brain, we combined the Cre/lox and the Tet system so that transgene expression was dependent on both, doxycycline induction and Cre recombinase. The general principle is depicted in Figure 1a. Quintuple (5x) genetic engineering of brain tissue with transgenic mice and/or adeno-associated viruses (AAV) provides rapid manipulation via the floxed, two-component Tet system (2x), cell type-specificity mediated by Cre recombinase (1x), prior labeling of cells primed for tet-dependent expression (1x, here: transgenic floxed tdTomato), and optical analyses of the induced genetic interference (1x, here: viral GCaMP6f). Cre recombinase can either be provided through transgenic mice, for example Parvalbumin-Cre mice, or through injection of a virus, for example Synapsin-Cre. The exciting strength of the viral approach is that the concentration of virus can be titrated according to the experimental needs. If only very few cells or even a single cell are intended to be manipulated, the Cre virus could simply be diluted considerably to transduce the approximate cell number. Conversely, for interference with many cells a higher titer is used. Thus, for all different Cre virus dilutions, the same high titer of the two Tet-On viruses can be used to achieve efficient transgene expression. As for the experimental time course, we typically waited 3-4 weeks after surgery for full transgene expression and surgery-related inflammation to cease (Xu et al., 2007). Prior to doxycycline administration, extensive baseline recordings (activity, morphology, etc.) of the same tdTomato+ cells and their surrounding cells were performed for a few days by means of *in vivo* optical imaging. Following tet-dependent transgene induction, the same cells and cellular parameters were repeatedly imaged in a longitudinal experimental design ranging from hours to days and potentially weeks. The unique approach of extensive imaging before, during, and after transgene induction of specifically tailored fluorescent readouts we term ‘optical interrogation’.

**Figure 1:**
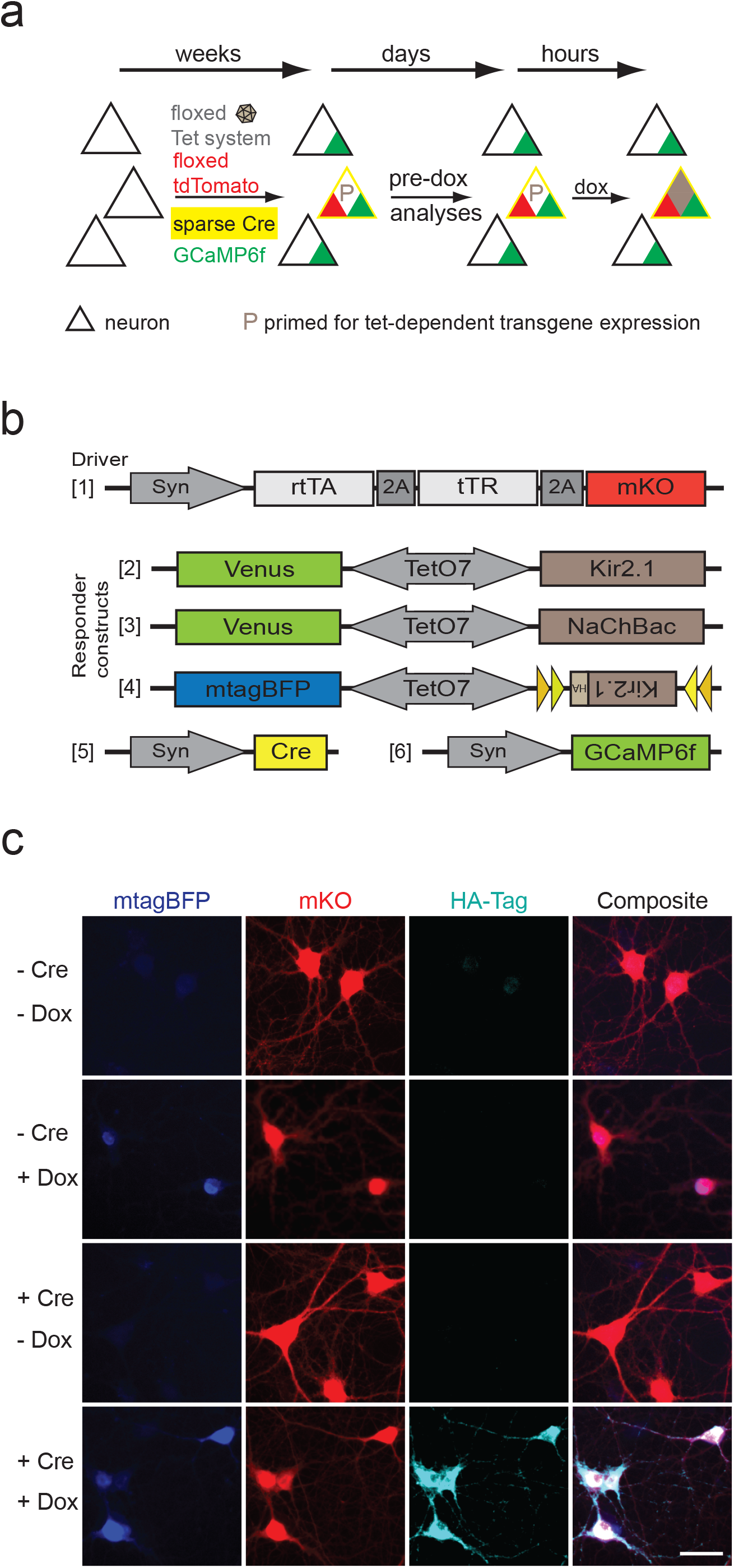
Establishment of a Cre-dependent Tet system. (a) Scheme and timeline for Cre- and tet-dependent manipulation of cells *in vitro* and *in vivo*. Co-expression of a Cre-dependent fluorescent marker (tdTomato) marks those cells that are primed for fast, tet-dependent expression of the desired transgene. (b) The optimized driver AAV containing reverse tetracycline-dependent transcriptional activator (rtTA) and tetracycline-dependent repressor (tTR) was used with either of three different responder AAVs. (c) In hippocampal cell culture, neurons were transduced with AAVs [1] and [4]. For cells labeled with ‘+Cre’, viral Synapsin-Cre [5] was co-administered. Expression of blue fluorescent mtagBFP required doxycycline for induction while immunodetection of the HA-tag required doxycycline and Cre recombinase (scale bar: 20 μm).

For the two component Tet-On system, we produced a ‘driver’ virus with the reverse tetracycline-dependent transcriptional activator (rtTA) [1] (Figure 1b) and ‘responder’ viruses containing the TetO7 promoter with the transgene [2-4]. We flanked the Kir2.1 transgene of one responder virus [4] with Cre recombinase recognition sequences while the mtagBFP transgene on the other side of the bidirectional TetO7 promoter was not modified. In dissociated hippocampal cultures, the dependency of the Kir2.1 transgene on doxycycline and Cre recombinase could be demonstrated using AAVs [1,4,5]. Doxycycline administration alone induced blue mtagBFP fluorescence, while the presence of both, Cre recombinase and doxycycline, was necessary for Kir2.1 expression (Figure 1c).

For genetic manipulation of neuronal activity, we chose the inward rectifying potassium channel Kir2.1 (Burrone et al., 2002) (neuronal hyperpolarization, eventually silencing the neuron) and the bacterially derived sodium channel NaChBac (Lin et al., 2010) (neuronal depolarization, eventually activating the neuron). Both transgenes were first tested in hippocampal cultures *in vitro*. Overexpression of Kir2.1 in neuronal cultures correctly targeted the exogenous Kir2.1 channel to the soma and dendritic compartments. As was previously shown, Kir2.1 was also localized in dendritic spines (Figure 2a) (Howe et al., 2008). In these cultures, HA-tagged Kir2.1 could be detected by immunofluorescence up to at least 96 hours after a 4 hour pulse of doxycycline (Figure 2b). This suggested that a single induction provides Kir2.1 protein for several days and that silencing via the Tet system can be long-lasting. To mimic the *in vivo* situation, analysis of tet-dependent manipulation of neuronal activity *in vitro* was performed by recording calcium transients via imaging of GCaMP6f fluorescence. Doxycycline application and induction of transgenes lead to a rapid and significant decrease (Kir2.1) or increase (NaChBac) of neuronal activity within hours (Figure 2c). In fact, initial trends of changing activities, albeit not significant, were observed after one hour. Together, these results suggest that the Tet system is suitable for rapid genetic manipulation of neuronal activity.

**Figure 2:**
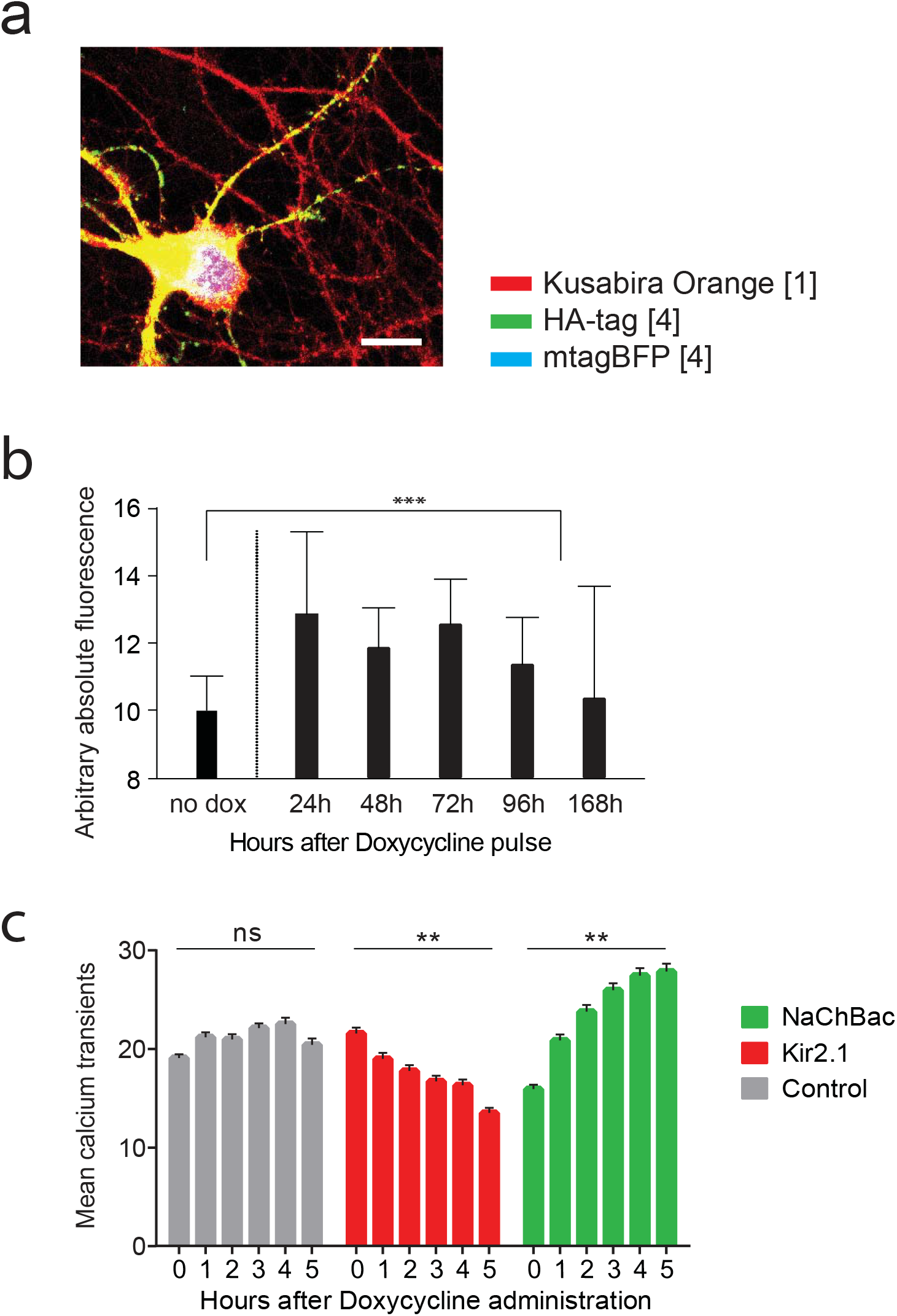
Genetic manipulation of neuronal activity in dissociated hippocampal cultures. (a) Overexpressed Kir2.1 was detected in the soma and dendrites, including spines. (b) A 4-hour pulse of 10 μM doxycycline produced long-lasting Kir2.1 protein presence *in vitro*. Neuronal cultures were transduced with the standard driver AAV [1] and an AAV containing a tet-dependent HA-tagged Kir2.1 construct [4]. Cultures were fixed, immunostained for the HA-tag at the indicated times, and the immunofluorescence quantified using NIH Fiji. At least 60 cells from two different cultures were quantified per time point (Wilcoxon signed-rank test, P< 0.0001; error bars: SD). (c) Tet-dependent induction of Kir2.1 [AAVs 1,2,6] or NaChBac [AAVs 1,3,6] significantly decreased (P=0.0011) or increased (P=0.0035) calcium signals recorded with GCaMP6f, respectively (≥ 4 hippocampal cultures and > 400 neurons per condition; statistical test: linear regression fit; error bars: SD).

Our early attempts to establish an efficient method for inducible transgene expression based on the Tet-On system were often flawed by considerable background expression in the absence of doxycycline (data not shown). This ‘leak’ expression from the TetO7 promoter is likely due to the presence of a minimal CMV promoter in the construct. This element cannot be removed because it is necessary for transcription. Thus, we decided to increase tightness via the driver virus by introducing an additional tetracycline-dependent repressor (Deuschle et al., 1995) (tTR). In the absence of doxycycline, the repressor binds to the TetO7 promoter thereby blocking unspecific ‘leak’ expression, but dissociates from DNA upon binding to tetracyclines. In cell lines, this approach has been shown to reduce leak expression at low doxycycline concentrations (Freundlieb et al., 1999). In dissociated hippocampal neurons, we found that the addition of tTR decreased the effective concentration (EC50) 3-fold (3.3 μM with tTR, 9.3 μM without tTR) (Figure 3a; Suppl. Figures 1a,b). While we expected a higher EC50 value with tTR and not without, this finding does not impact the effectiveness our method. Moreover, compared to the more gradual increase detected without tTR, its presence produced a step-like dose-response curve which should also improve tightness of the system. To test if the *in vitro* findings could be translated to *in vivo* experiments, we compared different virus combinations for their effectiveness in blocking fluorescent Venus ‘leak’ expression from the TetO7 promoter in the absence of doxycycline. In the living brain, the ‘leak’ expression could be successfully blocked by the presence of tTR. Within the same brain, a side-by-side comparison between a virus pair with tTR (AAVs 1,2) and a pair without tTR (AAV driver with luciferase instead of tTR + AAV 2) clearly showed that Venus expression was not apparent on the right hemisphere with the standard driver containing tTR (Figure 3b, top). To clearly differentiate green Venus signal from the pervasive green background fluorescence of the tissue, the blue channel of the same RGB image is shown below with the minor blue Venus fluorescence readily visible only in the left hemisphere (Figure 3b, bottom). To address the possibility that the leak expression of the TetO7 promoter stems from residual rtTA activity in the absence of doxycycline, we injected one hemisphere only with the Venus responder virus and no driver. Similar to the previous experiment with the tTR-less driver, abundant Venus expression was observed (Figure 3c). Again, the control hemisphere with AAVs [1,2] exhibited negligible Venus fluorescence. We conclude that tTR is a critical component for the Tet system *in vivo*, essentially limiting leak expression from the TetO7 promoter.

**Figure 3:**
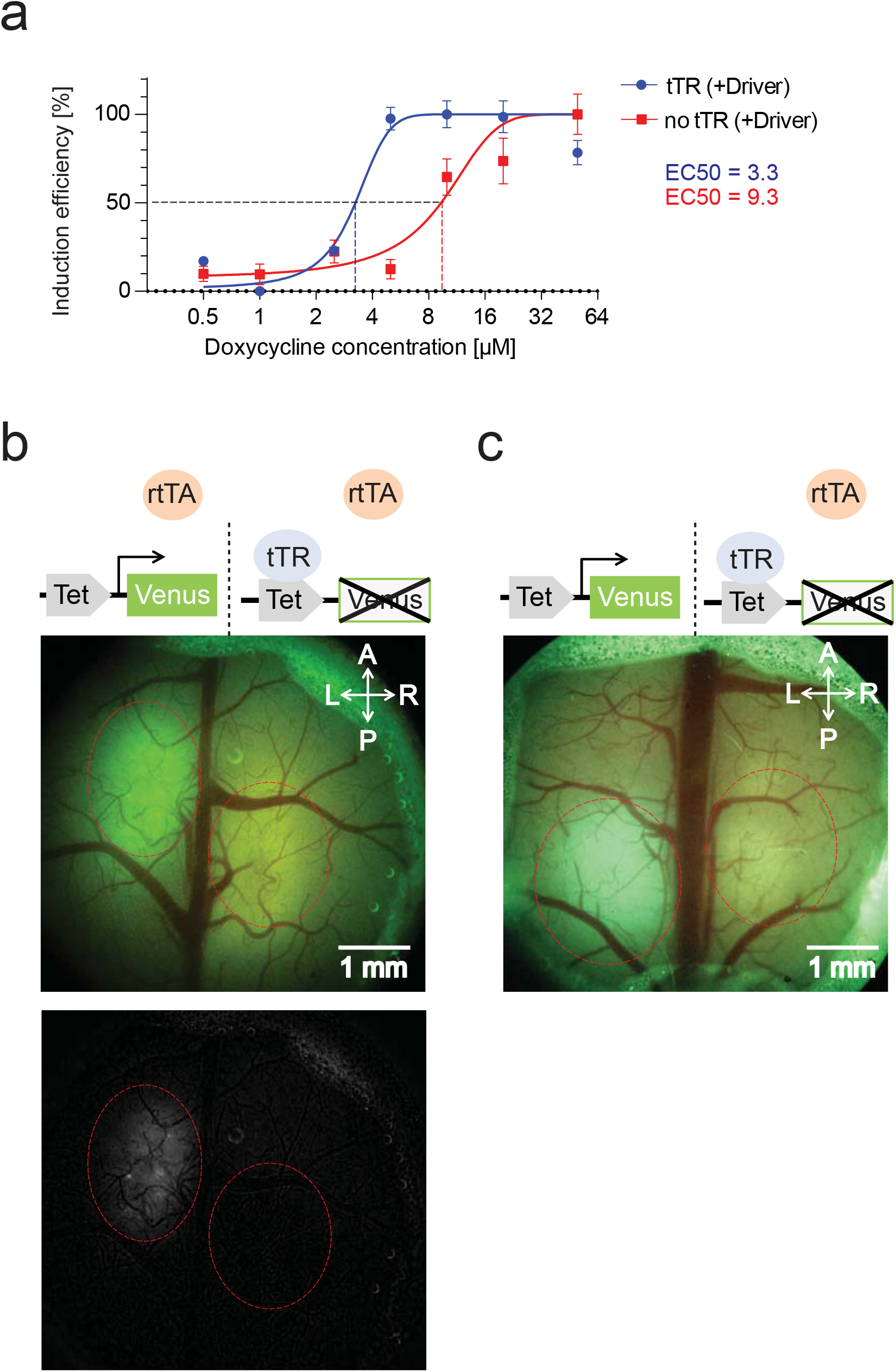
The presence of the tTR repressor affected tet-dependent Venus expression *in vitro* and *in vivo*. (a) Transduction of dissociated hippocampal cultures with [2] plus driver AAV [1] containing rtTA and tTR (blue) produced a different doxycycline dose-response curve than a driver AAV containing only rtTA and luciferase (red). Quantification of Venus fluorescence and curve-fitting computed two distinct EC50 values (on average 33 neurons quantified per condition). (b) *In vivo* leak expression in the absence of doxycycline was reduced by the presence of the repressor tTR. Top: Different virus combinations were injected into either hemisphere of a mouse brain. Green Venus leak expression was apparent in the left hemisphere without tTR. On the right, the presence of tTR substantially reduced Venus fluorescence. Stippled lines roughly outline the injection areas. Note the considerable green background fluorescence of brain tissue and skull. Bottom: Blue channel only display of the top image highlights minor blue Venus fluorescence on the left, but not the right hemisphere. (c) Leak expression was due to the Tet promoter and not residual rtTA activity because the responder virus alone produced Venus fluorescence on the left.

To achieve rapid and robust onset of doxycycline-induced transgene expression *in vivo*, we first compared two different routes of doxycycline administration, i.e. intraperitoneal injection vs. direct injection into the brain. For direct injections, we implanted chronic window glass coverslips with an acentric, re-sealable hole (Roome and Kuhn, 2014) of about ~ 0.5 mm (Figure 4a). This allowed repeated access to the brain so that doxycycline could be directly injected at any time after the surgery and window implantation. We also assessed transgene expression in hemispheres ipsilateral vs. contralateral to the injected hemisphere. In longitudinal two-photon microscopy experiments, for each mouse the levels of Venus fluorescence were repeatedly monitored in the same imaging plane. Imaging for all *in vivo* experiments typically occurred at a depth of about 150-250 μm below the brain surface in layer 2/3 of M1 or S1 depending on the injection site. Following induction, intense Venus fluorescence was observed by two-photon microscopy (Figure 4b). Note that the Venus fluorescence prior to doxycycline treatment was negligible. More importantly, because our approach allows analysis of the same cells before and after transgene induction, those very few cells with ‘leak’ expression could potentially be excluded in the post-hoc image processing, if necessary. In general, we found that transgene expression ipsilateral to the injection was substantially higher than in the contralateral hemisphere or after intraperitoneal injection (Figure 4c; Suppl. Figures 2a,b). We expected that injection into one hemisphere would lead to comparable transgene induction across the brain but probably doxycycline is cleared rapidly. The significant expression increase with direct vs. intraperitoneal injection is presumed to be a consequence of doxycycline not having to cross the blood-brain barrier (Beard et al., 2006) so that higher concentrations can be achieved in the brain. Thus, for all subsequent GCaMP6f-based (Chen et al., 2013) calcium imaging experiments, doxycycline induced silencing of neurons was assessed ipsilateral to the injection. Overall, we consider the levels of transgene expression that can be achieved with our approach *in vivo* after a single induction of doxycycline more than adequate for typical experimental needs.

**Figure 4:**
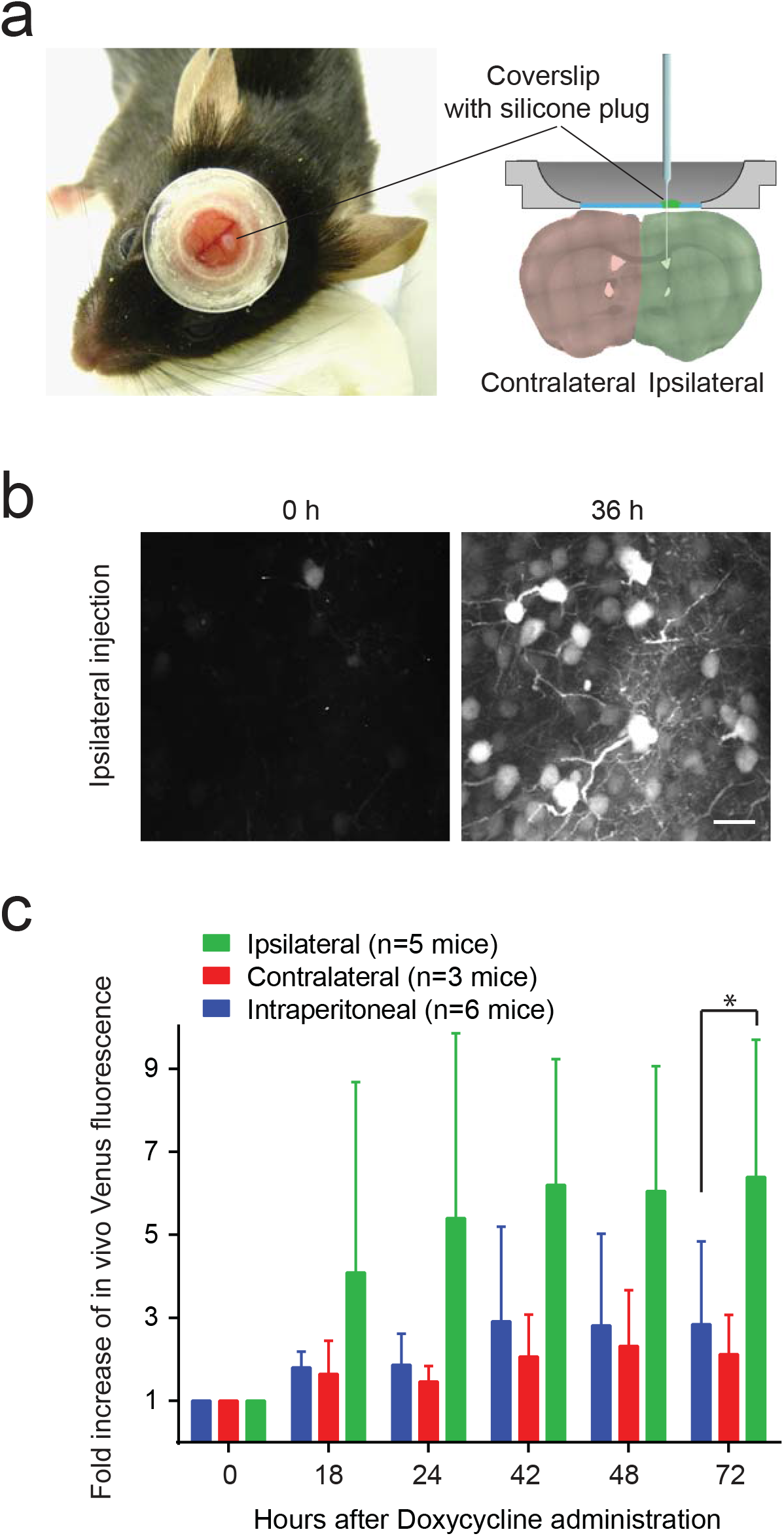
*In vivo* induction of tet-dependent transgenes by direct doxycycline injection into the brain. (a) Direct injection of doxycycline through a small hole in the chronic window. The hole could be resealed with silicone and thus allowed immediate and continued access to the brain. Imaging was possible ipsi- or contralateral to the injection site. (b) *In vivo* induction of Venus [AAVs 1,2] and two-photon imaging of the same S1 cortical region before and after doxycycline administration (scale bar: 20 μm). (c) Significantly increased ipsilateral (green) Venus *in vivo* expression compared to intraperitoneal (i.p.) injection (blue) [AAVs 1,2] (Kolmogorov-Smirnov test, P=0.026; error bars: SD). Expression contralateral (red) to the doxycycline injection produced transgene levels similar to i.p. injection.

In a first set of experiments to genetically manipulate neuronal activity *in vivo*, the population of neurons in which Kir2.1 will be induced by doxycycline was defined by the co-transduction with a dilute Synapsin-Cre virus [5] and the presence of Cre recombinase. A quadruple cocktail of viruses (driver [1], floxed Kir2.1 responder [4], dilute Synapsin-Cre [5], Synapsin-GCaMP6f [6]) was injected into transgenic Ai14 mice containing a floxed tdTomato reporter (Madisen et al., 2010) (Figure 5a). The contralateral hemisphere only received the GCaMP6f virus but not the two Tet-On viruses and served as control. Sparse, neuronal Cre expression (Cre+) could be visualized by tdTomato fluorescence which in turn identified those cells that were also primed for silencing via Kir2.1 and the Tet system. Because the Cre and GCaMP6f virus were both driven by the Synapsin promoter, the tdTomato signal typically colocalized with the green GCaMP6f fluorescence (Figure 5a). Because of the higher titer of the GCaMP6f virus, many additional neurons only exhibited GCaMP6f fluorescence but were negative for Cre recombinase (Cre-); the activity of these surrounding control neurons was assessed in parallel. Thus, in longitudinal imaging experiments using mildly anesthetized animals, spontaneous neuronal activity was repeatedly monitored over time by GCaMP6f fluorescence in the same neurons, with and without tdTomato signal. Mild anesthesia allowed analyzing relative changes of spontaneous activity and the anesthesia depth was continuously adjusted to a breathing rate of about 120 breaths per minute (Ellwardt et al., 2018) to ensure comparable activities across the longitudinal experiments. Because both the spontaneous activity and the breathing rates co-vary with the depth of anesthesia (Suppl. Figure 3a), the breathing rate was used as a proxy for the spontaneous activity. The breathing rate could be acutely monitored with a piezo element placed under the mouse thorax and the levels of isoflurane were adjusted manually if necessary. This ensured constant breathing rates during and across imaging sessions (Suppl. Figure 3b). Otherwise, we found that the variability of spontaneous activities was much more pronounced than the experimentally induced changes (data not shown). In addition, great care was taken to record the same cells in the same imaging plane despite returning the animals to their home cages in between time points. Each time point thus consisted of at least several dozen cells in the imaging plane whose spontaneous GCaMP6f activity could be recorded during a 9 minute movie. We established a semi-automatic pipeline to be able to extract and quantify these neuronal activities in longitudinal experiments. Briefly, maximum intensity projections (MIP) of all 10-12 imaging planes (i.e. time points) from each animal were first aligned and registered so that all the cells in the planes could be superimposed (Suppl. Figures 4a,b). Remarkably, for some experiments a near perfect superimposition was achieved (Suppl. Figure 4c). By concatenating the corresponding movies, the activity of individual neurons could be viewed and regions of interest (ROI) around the somata were manually determined for all recordable cells. To exclude ‘contaminating’ neuropil signal from our analyses, a second, doughnut-shaped ROI surrounding each cell was digitally created and subtracted from the somata ROI (Suppl. Figures 5a,b). Finally, shifting baselines of the concatenated GCaMP6f calcium transients were corrected with a customized MatLab curve fitting toolbox (Mathworks) (Suppl. Figure 5c). We then used the MLspike algorithm to extract and quantify action potential spiking from calcium transients (Deneux et al., 2016). A single dose of doxycycline induced rapid Kir2.1-dependent silencing in tdTomato-positive (Cre+) neurons within hours, while tdTomato-negative neurons remained unaffected (two-sided ANOVA Cre+ vs. Cre-, P<0.0001) (Figure 5b). Similar to the *in vitro* detection of Kir2.1 for up to 96 hours, reduced spontaneous activity persisted *in vivo* for at least 80 hours. Overall we find that most analyzed Cre+ neurons had decreased spontaneous activity while a minor fraction exhibited an increase (Suppl. Figures 6a,b). This is in stark contrast to Cre-neurons for which about the same number of neurons showed an increase or decrease of spontaneous activity, respectively (Suppl. Figures 6c,d).

**Figure 5:**
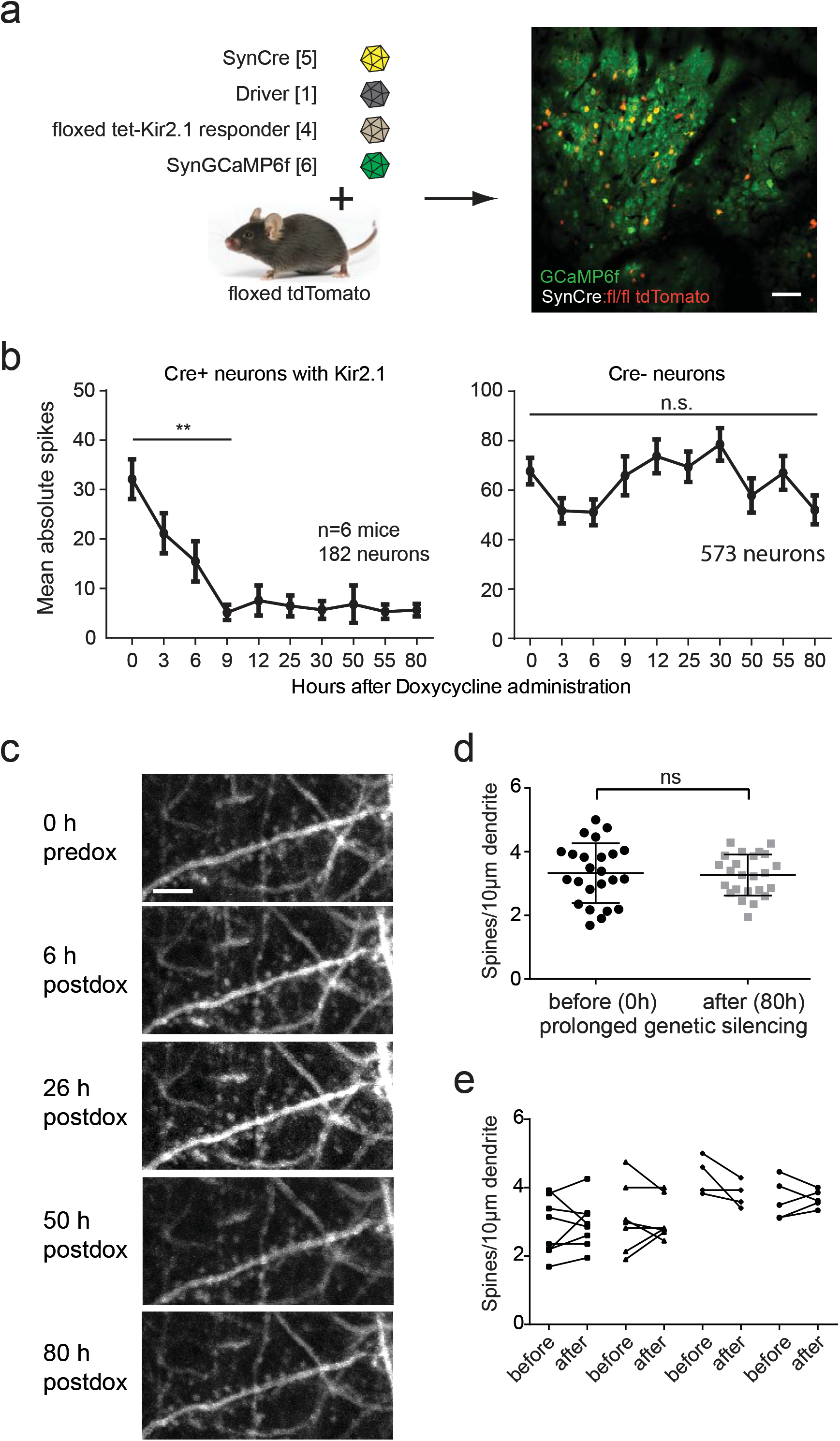
Rapid and long-lasting *in vivo* genetic silencing by induction of Kir2.1 in a sparse population of cortical neurons. (a) Genetic engineering of brain tissue to allow Cre- and tet-dependent transgene expression *in vivo*. Single *in vivo* imaging plane shows that yellow Cre+ neurons (red tdTomato + green GCaMP6f) can be clearly distinguished from green Cre-neurons (GCaMP6f only) (scale bar: 100 μm) (b) *In vivo* two-photon imaging of GCaMP6f: spontaneous cortical activity in Cre+ neurons (left) significantly decreased after tet-dependent induction of Kir2.1 compared to baseline (0 hours), while control Cre-neurons on the same hemisphere without Kir2.1 did not change (right). Displayed is the mean spiking of all cells (6 mice, ~30 neurons for each mouse; linear regression analysis: Cre+ [0-9 hours] P=0.0066, Cre- [0-9 hours] n.s., [0-80 hours] n.s.; error bars: SEM) (c) Representative example of longitudinal imaging of tdTomato fluorescence to assess morphological changes of manipulated neurons over time (scale bar: 5 μm). (d) Quantification of spines on dendritic stretches of neurons (c) that were silenced with Kir2.1 revealed no changes in spine numbers after prolonged silencing of more than three days (4 mice, 24 dendritic stretches, statistical test: nonparametric Wilcoxon matched-pairs signed rank test, P=0.4774). (e) Each before-after pair represents one dendritic stretch before doxycycline injection and 80 hours after Kir2.1 induction. Dendritic stretches from each mouse were grouped together. No statistical difference was observed for either mouse (statistical test: nonparametric Wilcoxon matched-pairs signed rank test; from left to right: P=0.7813, P=0.6250, P=0.250, P>0.999)

**Figure 6:**
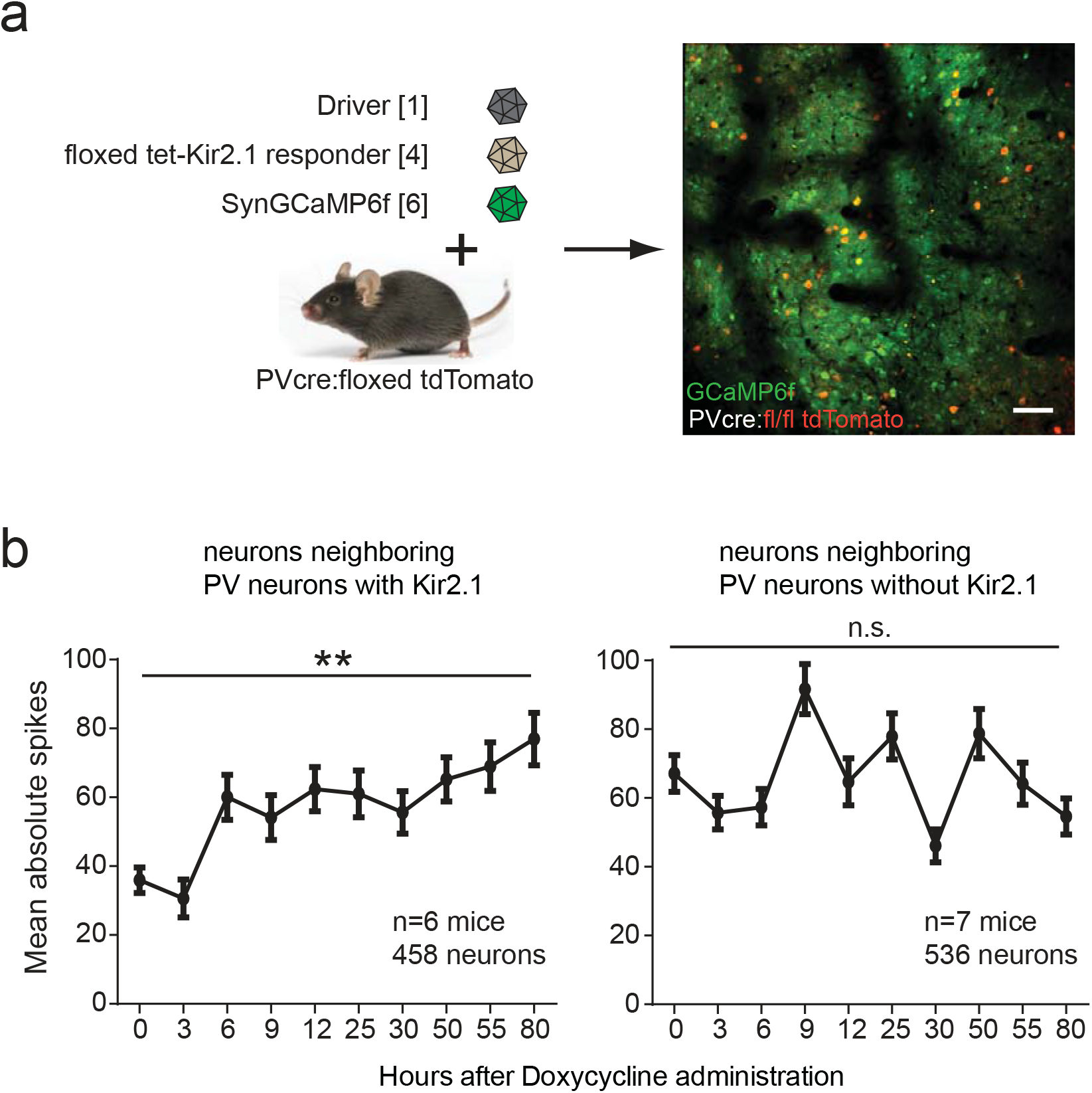
Cell type-specific silencing of inhibitory parvalbumin neurons increased neuronal activity of surrounding cells. (a) Genetic engineering of brain tissue to allow tet-dependent silencing of parvalbumin neurons. Single *in vivo* imaging plane shows that yellow parvalbumin neurons (red tdTomato + GCaMP6f) can be clearly distinguished from the surrounding green Cre-neurons (GCaMP6f only). (b) Silencing of parvalbumin neurons significantly increased neuronal activity of surrounding cells (left). Neurons in control hemispheres of the same mice but without Kir2.1 did not show any significant change in activity (right). Displayed is the mean spiking of all cells (6/7 mice [due to technical reasons, the experimental hemisphere of one mouse could not be recorded], ~50-100 neurons for each mouse; linear regression analysis: Cre+ P=0.0073, Cre-n.s.; error bars: SEM).

Burrone and co-workers previously demonstrated in neuronal cultures that Kir2.1-mediated silencing after synapse formation did not lead to homeostatic changes in spine number (Burrone et al., 2002). So far, these findings were not recapitulated *in vivo*. We therefore used dendritic tdTomato fluorescence for longitudinal analysis of neuronal morphology in these silenced neurons (Figure 5c). Notably, the same Cre+ neurons and their dendritic spines could be repeatedly reidentified in consecutive imaging sessions which permitted quantification of spine density changes. No significant spine changes could be detected 80 hours after prolonged reduction of neuronal activity (Figure 5d). This overall trend of spine stability was also seen when analyzing individual dendritic stretches from four different mice as there were only small changes observed (Figure 5e). Our data thus extend recent findings that showed synapse formation during development in the absence of neuronal activity (Sando et al., 2017; Sigler et al., 2017) to synapse maintenance despite acute, prolonged silencing in adult mice.

The use of a diluted Synapsin-Cre virus afforded great flexibility in the number of transduced Cre+ cells with our method. Alternatively, the abundance of transgenic Cre mice offers great flexibility for cell type-specific manipulation with our method. Taking advantage of such mice, we wanted to demonstrate the cell type-specificity by silencing inhibitory parvalbumin (PV) neurons in PV-cre mice *in vivo*. The prediction was that in this set of experiments, the surrounding non-PV cells change their activity because of the importance of PV cells in regulating neuronal activity and homeostasis. A quadruple virus cocktail ([1],[4],[6] + [CAG-floxed GCaMP6f] CAG: CMV early enhancer/chicken beta actin promoter) was injected into double transgenic parvalbumin-Cre, Ai14 mice. Red tdTomato fluorescence clearly differentiated PV neurons from non-PV cells (Figure 6a). Silencing inhibitory PV-cells led to rapid and significant increases in spontaneous activity of surrounding non-PV neurons (two-sided ANOVA Kir2.1 vs. control hemisphere, P=0.0017) (Figure 6b). The control hemisphere was only injected with GCaMP6f and the corresponding changes of the calcium transients over time were not significant. While the biologically variability was expectedly high, in one animal the increase of spontaneous activity was up to ten-fold (Suppl. Figure 8a). Overall, the effect of silencing PV neurons persisted in non-PV cells for at least 80 hours as their activity remained high. We also attempted to characterize changes in PV activity after induction of Kir2.1. However, since the synapsin promoter preferentially labels excitatory neurons, only a fraction of PV cells showed GCaMP6f fluorescence (Suppl. Figure 7). Therefore, we co-injected a CAG-floxed GCaMP6f virus to monitor the activity of PV neurons but presumably their spontaneous calcium transients were too frequent or minor for quantification (data not shown). Because the observed changes in surrounding neurons temporally correlated with the Kir2.1 induction, were of increasing nature as expected from silencing inhibitory neurons, and were based on a well-characterized PV-cre mouse line, we conclude that PV neurons were specifically and acutely silenced *in vivo*. We next wondered if there was also a spatial correlation between the induced manipulation of Cre+ neurons and the observed changes in the surrounding Cre- cells. In a neuronal network within the imaging plane, differences in the local connectivity are highly likely because of cell distances and cell types. Depending on the extent of this connectivity, cells may exhibit different changes downstream of the induced genetic manipulation. We therefore analyzed the changes in spontaneous activity in non-PV neurons as a function of average distance to PV neurons 80 hours after induction. Within the imaging plane (Figure 7a), tdTomato+ and GCaMP6f+ cells were digitally identified, catalogued, and the computed activity change of GCaMP6f+ cells assigned. The degree of activity change was highlighted as a heat map of the imaging plane with PV cells shown in black (Figure 7b). We found that non-PV cells which had a smaller average distance to all detected PV neurons were significantly more affected (Figure 7c). The observed maximum ‘functional reach’ of PV cells of about 500 μm (intersection with x-axis) is in good agreement with previously published morphological characteristics of PV neurons suggesting a ‘morphological reach’ of 760 μm +/−130 μm (Halasy et al., 1996). For comparison, no distance dependency was seen between PV cells and the surrounding neurons in control hemispheres that were only injected with GCaMP6f, but without driver and responder (Figure 7d). We also repeated this type of analysis for the Synapsin-Cre injected mice. While the surrounding Cre- neurons as a population did not display any change in activity (Figure 5b), possibly small changes of nearby Cre- neurons could have been missed. Within the imaging plane (Figure 7e), a heat map of activity changes for Cre+ (black) and Cre- neurons was generated (Figure 7f). However, no distance-dependent changes could be observed for either neuron population (Figures 7g,h). Taken together, the data suggest that induced changes in activity in one neuronal population can have secondary effects on surrounding neurons and that these secondary activity changes may be a function of distance and connectivity.

**Figure 7:**
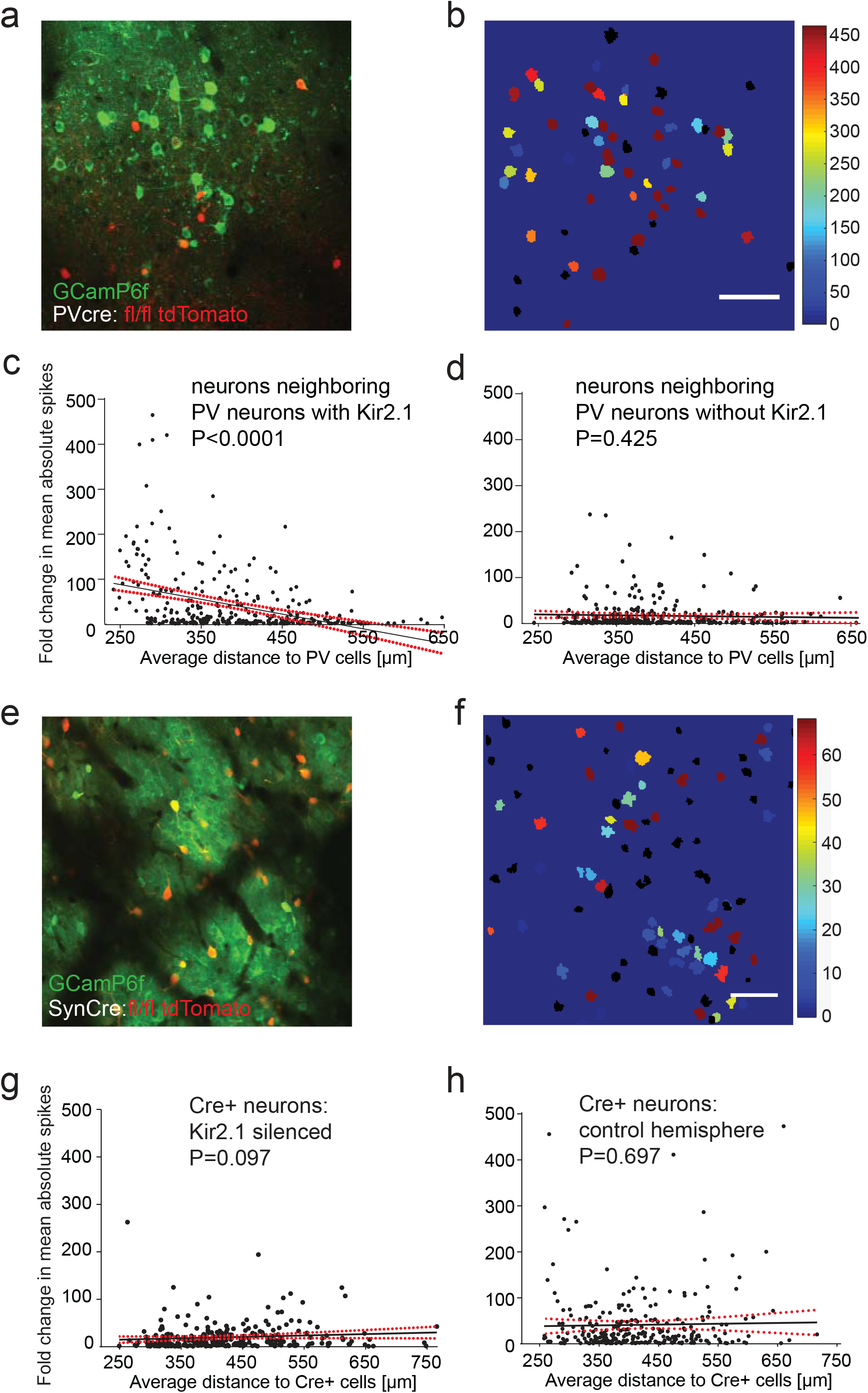
Silencing a neuronal subpopulation *in vivo* and its distance-dependent effect on the activity of surrounding neurons. (a) *In vivo* image of a cortical region showing red/yellow PV cells and green fluorescent surrounding neurons (GCamP6f) (b) Heat map representation of the same region as in (a). The changes in average spiking relative to the initial activity over all recordings were color coded. The positions of PV cells are displayed in black (scale bar: 100 μm). (c) Each dot represents one neuron displaying its average distance to all detected nearby parvalbumin neurons and its change in neuronal firing. Surrounding neurons with a shorter average distance to PV cells displayed significantly higher increases in activity compared to those further away (linear regression analysis: P<0.001). (d) Control neurons on the other hemisphere with PV cells which did not express Kir2.1 showed no distance-dependent changes in neuronal activity (linear regression analysis: n.s.). (e) *In vivo* image of a cortical region showing red/yellow Cre+ cells and green fluorescent surrounding neurons (GCamP6f) (f) Heat map representation of the same region as in (e). The changes in average spiking relative to the initial activity over all recordings were color coded. The positions of Cre+ cells are displayed in black (scale bar: 100 μm). (g) Each dot represents one neuron displaying its average distance to all detected nearby Cre+ cells and its change in neuronal firing. Injection of a dilute Synapsin-Cre AAV defined the Cre+/tdTomato-positive neuronal subpopulation which was silenced with Kir2.1. The average distance of manipulated neurons to GCamP6f-only neurons had no significant effect on the observed changes in activity. (h) Control hemispheres in the same animal were only injected with the dilute Synapsin-Cre and the Synapsin-GCamP6f AAV, but not the two driver and responder AAVs of the Tet system. No distance-dependent change in activity was detected.

## Discussion

Our method provides **high**-resolution, inducible interference and interval imaging of individual cells (**high I^5^**) which is why we designate it the ‘HighFive’ approach. The ‘HighFive’ approach proved to be versatile, robust, and is readily implemented in longitudinal *in vivo* imaging paradigms. Prior marking of cells with a Cre-dependent, fluorescent reporter accurately predicts when and where Tet-dependent transgene expression occurs, thereby allowing precisely tailored *in vivo* optical interrogation and manipulation of defined cell types. This is currently not possible with any other conditional expression method and opens up new possibilities for cellular and network-level neuroscience research. For example, we envision improved rescue experiments by temporally controlled, cell type-specific expression of the relevant transgene in genetic knock-out animals. With the HighFive approach, transgene expression can exclusively be correlated in time and space to potentially subtle, cellular phenotypic changes - within the same cell, but also within the context of the cellular network *in vivo*. Thus, our method offers exciting new possibilities to study cell biology *in vivo* at the level of single cells.

Establishing stable and reproducible experimental conditions *in vitro* is usually straightforward, it is however a big problem in living animals, for example when studying neuronal mechanisms of information processing. Consequently, characterizing acute genetic manipulations *in vivo* poses a tremendous technical and logistical challenge: matching control and experimental animals to have the same level and time course of expression, same age, gender, depth of anesthesia, imaging quality, amongst other factors. Here, instead of matching two different cohorts of animals (experiment vs. control) and requiring larger animal numbers, our longitudinal approach allows comparisons before and after transgene expression in the same cells and animals. The effects of transgene expression can be assessed in one extended imaging session, or as we showed, in longitudinal experiments.

In addition, by combining the Tet with the Cre/lox system, our method offers both, rapid transgene expression conferred by the Tet system and exquisite cellular specificity defined by the Cre/lox system. Gene expression, from DNA to protein, typically happens in less than one hour (Shamir et al., 2016). Rapid, inducible transgene expression via the Tet-On system can approximately recapitulate this time course (Hasan et al., 2001). We are, however, not aware of any publication that demonstrated similarly fast expression times for the Tamoxifen inducible Cre/lox system. In vitro and *in vivo* experiments suggest instead that the combination of (4-OH) Tamoxifen pharmacokinetics, Cre-mediated DNA recombination, and transgene induction appear to require at least several hours for detectable expression (Fossat et al., 2006; Hayashi and McMahon, 2002). Even longer time courses were reported for the Tet-Off system (Kistner et al., 1996). The Tet system can provide high spatial resolution (Cambridge et al., 2009) and cellular specificity using transgenic mice (Tillack et al., 2015) or viruses (Watakabe et al., 2017), but for both, the Tet-On and Tet-Off systems, there are only about two dozen mouse lines available at Jackson Labs with neuron specific promoters. Conversely, over six hundred Cre recombinase mouse lines have been created for targeted genetic manipulation of neurons of specific cell types and in specific brain regions (Tsien, 2016). The HighFive method takes advantage of the cellular specificity provided by the many available Cre mouse lines particularly since both, standard Cre lines and inducible CreERT2 lines can be used as long as Cre recombinase is active before doxycycline induction. We also improved the Tet-On system for *in vivo* work by introducing the tTR repressor for less background expression and by direct brain injections for higher inducibility compared to intraperitoneal injections.

The second major methodological advancement is the development of a new approach for conditional manipulation of neuronal activity. We used the cellular precision of the ‘HighFive’ method to successfully interfere with neuronal activity in defined neuronal populations. The high temporal resolution of genetic transgene expression allowed acute manipulation of neuronal networks with the experimental intention of being able to study network homeostasis *in vivo*.

To deliver a proof-of-principle experiment, we silenced a subpopulation of mostly excitatory neurons as defined by Synapsin-Cre expression, but did not observe any obvious homeostatic compensatory mechanisms. The silenced neurons did not exhibit significant morphological changes while the activity of the surrounding neurons also appeared unaffected. *In vitro*, Kir2.1 expression, after the bulk of synapse formation had occurred, also did not alter spine density. However, Kir2.1 expression in cultured neurons reduced their spiking frequencies which returned to control levels after three days (Burrone et al., 2002). To accommodate this finding, the authors proposed a homeostatic increase in synaptic input strength that we could not detect with the analysis method available to us, i.e. somatic GCaMP6f transients. Not surprisingly, different homeostatic mechanisms may thus operate *in vitro* vs. *in vivo*. Conversely, we find that silencing parvalbumin cells did increase the activity of surrounding neurons *in vivo*. Most likely this increase is simply a reflection of less inhibition by the parvalbumin neurons, but theoretically, a possible minor homeostatic effect cannot be excluded. In fact, we would like to speculate that the temporary reduction seen after the first rapid rise of activity (individual mouse, hours 30-55, Suppl. Figure 8a) could be a consequence of the network attempting to compensate the activity increase only to eventually succumb to the strong impact of the cumulative Kir2.1 expression. We did observe a similar decrease in activity in two other mice, albeit not as pronounced (average of three mice, Suppl. Figure 8b). Future work will have to investigate if this observation is reproducible and will have to aim at identifying possible underlying mechanisms.

In general, being able to silence or hyperactivate neuronal firing is central to many research questions. *In vivo*, we focused exclusively on genetic silencing via Kir2.1 because the interpretation of such ‘loss-of-function’ experiments is more straightforward. Silencing with the powerful optogenetics typically eliminates neuronal firing, but with our approach network activity was strongly reduced instead of completely abolished which might be more functionally relevant for some network experiments. Also, the genetic manipulation of neuronal activity via Kir2.1 or NaChBac represents a desirable complement to the DREADD system (Armbruster et al., 2007) whose active ligand was recently found to be the psychoactive compound clozapine (Gomez et al., 2017; MacLaren et al., 2016). Nevertheless, we do not see our method as competition to optogenetics or DREADDs but rather as an important, complementary approach which is perhaps better suited for behavioral experiments. Also, since our protocol only involves a single transgene induction that is effective for at least three days, this is particularly attractive for activity-dependent manipulation of behavior since acutely administered clozapine N-oxide (CNO) decreased to very low plasma levels within 2 hours (Guettier et al., 2009).

## Acknowledgments

We thank Gabriele Krämer, Michaela Kaiser, and Claudia Kocksch for technical assistance. Albrecht Stroh and Amit Agarwal provided valuable input to the manuscript. We are indebted to Thomas Kuner for continuous support of the project and comments on the manuscript. This work was funded in part by a doctoral fellowship of the Friedrich-Ebert-Stiftung (F.T.), a Frontiers grant (S.C.), and the SFB 1134 (S.C.)

## Author Contributions

S.C. conceived and supervised the study, and wrote the manuscript. F.T. developed and performed all the experimental work and data analysis. J.K. developed analytical tools and contributed substantially to imaging and data analysis. Both F.T. and J.K. developed the semi-automatic processing pipeline.

The authors declare no competing interests.

## Methods

### DNA constructs and plasmids

All plasmids were based on standard plasmid vectors with a rAAV backbone. A polycistronic vector of rtTA, luciferase, and mKO (Kusabira Orange) coupled via self-cleaving 2A peptide bridges and under control of the human synapsin promoter for neuronal expression (plasmid kindly provided by Rolf Sprengel) was used to clone the driver construct by exchanging luciferase with the repressor tTR. The ‘responder’ plasmids were all based on bidirectional TetO7 sequences flanked by a minimal Cytomegalovirus Immediate-Early (minCMV-IE) promoter (Gossen and Bujard, 1992). Viral DNA constructs [1]-[3] that were sufficiently below the total packaging limit of 5.2 kb had the mRNA stabilizing Woodchuck Hepatitis Virus Post-Transcriptional Regulatory Element (WPRE) inserted ahead of the Bovine Growth Hormone Poly-Adenylation Signal at 3’-end (Wang et al., 2016). For Cre-dependent expression, the Kir2.1 transgene was tagged with HA (hemagglutinin peptide tag) and flanked by DIO-sites (double-floxed inverse open reading frame) harboring LoxP & Lox2272 sites (Sohal et al., 2009).

### Recombinant adeno-associated viruses

For rAAV production a helper-free system was used that generates capsid chimeras of AAV serotype 1 and 2 at a ratio of 1:1. Briefly, 4×10^6^ HEK293T cells/cm² were plated in ten 15 cm Ø cell culture dishes and were grown in Dulbecco's Modified Eagle Medium containing 2x MEM NEAA (PAN Biotech), and Penicillin-Streptomycin (100 U/mL, Gibco). Transfections were performed 24 h after plating using the standard calcium phosphate method. DNA of the desired AAV and the two commercial helper plasmids (Plasmidfactory) pDP1rs and pDP2rs coding for the capsid proteins of the AAV1 and AAV2, respectively, were mixed at equimolar ratios to a final concentration of 37.5 μg DNA/plate. The DNA mixture was added to 6.8 mL of dH_2_O and 1 mL of 2.5 mol/L CaCl_2_ and vortexed briefly. 12 mL 2X HeBS buffer (280 mmol/L NaCl, 50 mmol/L HEPES, 1.5 Na2HPO4, adjusted to pH 7.05 with NaOH) was added dropwise while vortexing the CaCl_2_-DNA solution. After 100 s, 2 mL of the transfection mixture was added dropwise to each plate. 18 – 24 h later, the transfection efficiency should be at least 25 %. The medium was exchanged with 30 mL fresh DMEM and 48 h later, the cells were scraped from dishes and pooled with the supernatants. The suspension was centrifuged at 200 x g and 4 °C for 10 min. The supernatant was transferred to separate Falcon tubes and 125 U Benzonase per 50 mL supernatant were added and incubated for 2 h at 37C. The pellet was resuspended in TNT extraction buffer (20 mmol/L Tris-HCL pH 7.5, 150 mmol/L NaCl, 1 % Triton X-100, 10 mmol/L MgCl_2_) at room-temperature (RT) for 10 min to lyse the cell membranes. 1000 U of Benzonase were added and the solution incubated at 37 °C for 1 h. Cellular debris was removed by centrifugation (3600 x g at 4 °C, 15 min) followed by filtering of the supernatant with a 0.45 μm Millex PVDF membrane filter (Merck Millipore). For subsequent FPLC purification (Fast Protein Liquid Chromatography; ÄKTAprime plus, GE Healthcare) all solutions were degassed by vacuum-driven filtration (Stericup 500 mL Durapore 0.45 μm PVDF, Merck Millipore). Before loading, HiTrap Heparin HP (GE Healthcare) columns were first washed with 5 mL dH_2_O at 0.5 mL/min and then equilibrated with 5 mL PBS at 0.5 mL/min. The supernatant and the crude lysate were separately filtered using a 0.2 μm syringe Filter (Puradisc FP 30 Cellulose Acetate Syringe Filter, GE Healthcare) and directly loaded onto the column at 1 mL/min one after another. The column was then washed at a rate of 0.5 mL/min with 10 mL PBS and the virus eluted with elution buffer (500 mmol/L NaCl in PBS) at 0.5 mL/min. The eluates containing viral capsids were collected and transferred to an Amicon Ultra-15 filter tube (Merck Millipore). The filter tube was centrifuged at 3000 x g at 4 °C until less than 1 mL solution was left and the samples in the filter tube were subsequently washed twice with 13 mL PBS. The final run was continued until ~250 μL were left. This concentrated virus solution was filtered (0.22 μm Millex, Merck Millipore) and stored at 4 °C until further use. A simple qualitative assessment of the virus titer was done by transducing primary rat hippocampal cultures with 0.5 μL or 1 μL of virus and assessing fluorescence up to 7 days later. Viruses that did not produce satisfying fluorescence in cell culture were discarded and the preparation was repeated. For calcium imaging, a GCaMP6f rAAV (AAV1.Syn.GCaMP6f.WPRE.SV40) under the synapsin promoter was purchased from UPenn Vector Core (University of Pennsylvania, USA), diluted 1:10, and mixed with the driver/responder viruses 1:1:1. The Synapsin-Cre AAV was co-injected in dilution of about 1:500-1:2000 to achieve sparse transduction of neurons *in vivo*.

### Neuronal cell culture and in vitro assays

#### Neuronal Cell Cultures

Primary rat (Sprague-Dawley, Janvier) hippocampal neurons from P18.5 embryos were produced by a standard protocol (Banker and Goslin, 1998). The cells were plated in 24 well plates (Falcon Clear Flat Bottom TC-Treated Multiwell, Corning) at about 30.000 cells per well using 12 mm² poly-L-Lysine (Sigma-Aldrich) coated coverslips.

#### Immunofluorescence staining

Cultured neurons were transduced with rAAVs, induced with 10 μM doxycycline, and were subsequently fixated at defined time points by incubation in 1 mL 4 % Paraformaldehyde (Sigma-Aldrich) in PBS for 5 min. Immunofluorescence staining was performed by a standard protocol using an anti-HA rabbit primary antibody (C29F4, Cell Signaling Technology, diluted 1:1600) and a secondary antibody goat anti-rabbit Alexa Fluor 633 (Invitrogen, 1:1000 dilution). Analysis and quantification of epifluorescence images was achieved using Fiji (Schindelin et al., 2012).

#### *In vitro* calcium imaging

To study the effect of Kir2.1 or NaChBac on the activity of neurons *in vitro*, poly-L-Lysine coated cell culture dishes with a lid (μ-Dish 35mm, #81151, Ibidi, Martinsried) were used. Rat hippocampal cultures were transduced between DIV7 and DIV10 with 1 μL each of the Driver AAV, one of the Responder AAVs, and a 1:10 dilution of the commercial GCaMP6f AAV. After DIV15, cultures were continuously induced with a final concentration of 10 μM doxycycline and were continuously imaged for 5 minutes each hour for 5 hours. The imaging was performed on a Leica SP8 confocal microscope with a live cell imaging chamber at 37 °C. 5 % CO_2_ was superfused onto the medium with a cannula through the lid of dishes. For analysis, cell somata were manually selected by morphology independent of firing, calcium traces extracted, and the peaks quantified with a custom Matlab script.

#### Animals

All experiments were conducted in accordance with the German animal welfare guidelines and were approved by the Regierungspräsidium Karlsruhe. Mice were kept at a 12/12 – hour dark/light cycle synchronized with the local day-night cycle. Water and food were available ad libitum, except for the imaging sessions. Mice were generally housed in ventilated racks with up to 3 animals per cage, but were separated after surgery to minimize the risk of injury. Transgenic mice were either a homozygous Ai14 (B6.Cg-*Gt(ROSA)26Sor^tm14(CAG-tdTomato)Hze^*/J) mouse line with Cre-dependent tdTomato expression (JAX stock #007914 (Madisen et al., 2010)). Or, for cell-type specific manipulation of Parvalbumin (PV) interneurons, a double homozygous line derived from the Ai14 line and a PV-Cre (B6.129P2-*Pvalb^tm1(cre)Arbr^*/J) line (JAX stock #017320 (Hippenmeyer et al., 2005)).

#### Stereotactic injection, chronic cranial window implantation, and doxycycline administration

Surgery was performed on 8-12 week old mice. Injection of viruses and the implantation of a chronic cranial window were performed following craniectomy. The animals were anesthetized by an intraperitoneal (i.p.) injection of a mixture of 40 μl Fentanyl (1 mg/mL, Janssen-Cilag), 160 μL Midazolam (5 mg/mL, Hameln Pharma Plus) and 60 μL Medetomidin (1mg/mL, Sedan Alvetra-Werfft) at a dosage of 3.1 μL/g body weight. Bepanthen eye lotion (Bayer) was applied to both eyes to prevent dryness. To antagonize anesthesia at the end of the surgery, animals received a subcutaneous (s.c.) injection of a mixture of 120μL Naloxon (0.4 mg/mL, Inresa), 800 μL Flumazenil (0.1 mg/mL, Fresenius Kabi) and 60 μL Antipamezole (5 mg/mL, Pfizer) at a dosage of 6.2 μL/g body weight. The body temperature of the animals was maintained at 37-38 °C during the whole surgical procedure by a feedback-controlled heating pad (FHC). After the surgery was finished the animals received an s.c. injection of Carprofen (Rimadyl, Pfizer) dosed 10 μg/g body weight as an analgesic. The analgesic treatment was continued for at least two consecutive days or as long as the mice showed behavioral signs of pain(Langford et al., 2010). For the surgical procedure, the scalps of the mice were shaved and subsequently mounted in a stereotactic EM70G manipulator (Kopf Instruments). Prior to severing the skin at about 9 mm diameter around bregma, ~100 μL of 1 % Xylocain solution (Astra Zeneca) was applied subcutaneously, and the skin locally removed. After cleaning the skull surface, a 6 mm diameter coverslip (VWR) was used to mark a circular area with a pencil on the skull. This template was used to thin the skull at ~6.5 mm diameter using a hand-held dental drill (Osada EXL-40). The excised circular bone was removed by carefully lifting it up with blunt forceps. Potential bleeding was stopped with moistened coagulant foam (Equimedical) that contains fibrinogen. Subsequently, the dura was removed to improve long-term imaging results. Up to ~1.2 μL of viral solution per brain hemisphere were injected at cortical depths of 400 – 500 μm with a glass micropipette (Blaubrand IntraMARK 5 μl) pulled to about 10 – 20 μm inner diameter at the tips using a horizontal puller (Sutter Instruments). Usually 2–3 sites per hemisphere in S1 cortex were injected to increase the transduced brain area. The viruses were injected at a speed of about ~3 μL/h with manually controlled pressure applied using a 10 mL syringe. After positioning the injection micropipette at the desired depth, the tissue was allowed to tighten around the micropipette by waiting for about two minutes before starting the injection. Meanwhile, the brain surface was continuously kept moist with sterile phosphate buffered saline at all times to avoid drying out. When the injections were completed, the glass coverslip with a silicone filled access hole (Roome and Kuhn, 2014) (~0.9 mm diameter, custom made) was sterilized in 70 % Ethanol and chronically implanted. The access hole was placed above the ventricle (from bregma: Y= −0.7 mm, X= ± 1.5 mm) and the coverslip glued to the skull with dental acrylic cement (mixture of glue: Cyano [Hager & Werken, Duisburg] and powder: Paladur [Heraeus, Hanau]), while avoiding contact between cement and brain tissue. Additionally, the wounded skin around the surgery area was also sealed with dental cement. For animal mounting at the microscope setup, a round 3D-printed (Ultimaker 2, Geldermalsen, Netherlands) plastic crown was glued onto the skull. Following surgery, animals were treated with Dexamethasone (100 μL s.c. injection 1 mg/mL, Bela-Pharm) for two consecutive days to minimize bleeding and inflammation below the cranial window [surgery protocol adapted from(Holtmaat et al., 2009)]. Animals were allowed to recover from the surgical procedure and the inflammation reaction was allowed to cease for 3 to 10 weeks before experiments were conducted. For intracerebroventricular (ICV) injection of doxycycline, animals were anaesthetized with 1 % Isoflurane and mounted onto the stereotactic setup. The micropipette was leveled at glass surface and advanced slowly through the silicone access hole to 2.2 mm depth aiming for the ventricle (from bregma: X= ± 1.5 mm/ Y= −0.7 mm/ Z= −2.2 mm). 1.5 μL 8 mmol/L doxycycline Hyclate (Sigma-Aldrich, sterile filtered) in PBS were injected slowly (0.15 μl/min). To achieve maximal expression levels by a single induction protocol, 2.5 mg of 9TB doxycycline (Echelon) in 500 μL PBS were IP injected additionally to the ICV injections at the same time.

#### *In vivo* two-photon imaging

Mice were initially anaesthetized with 2 % Isoflurane (Baxter) and mounted onto the imaging stage. Before imaging, the cranial window was thoroughly cleaned with water and the volume above the cranial window was filled with distilled water. The site and expression of virus-mediated fluorescence was verified by epi-fluorescence microscopy coupled to a TriM Scope II multi-photon microscope (LaVision) using the imaging software Imspector of the same company. Imaging was performed with a 16X (NA=0.8) or 25X (NA=1.1) water immersion objective (both Nikon) using the appropriate filter sets (Venus or GCaMP6f: 535/70 nm and tdTomato: 650/100; Chroma). Fluorescence emission was detected with low-noise high-sensitivity photomultiplier tubes (PMTs, H7422-40-LV 5M; Hamamatsu). Two-photon imaging was performed at cortical depths between 80 and 130 μm and focal planes were chosen where many bright fluorescent layer 2/3 neurons were located. During imaging sessions, Isoflurane levels were continuously adjusted to about 0.8 to 1.5 % to yield respiratory rates between 110 and 130 per minute. Respiratory rates were monitored with a piezoelectric transducer (3 cm diameter, V/AC 1.3 ± 0.5 kHz, FT-31T-1.3A1-472) (Zehendner et al., 2013) and an amplifier from HEKA using a custom-written Matlab script. In this mildly anesthetized state, the animals exhibited substantial spontaneous cortical activity but no voluntary movement. For the respective panels in Figures 5, 6, and S8 the baseline recordings before doxycycline injection were averaged and grouped as time point ‘0 hours’.

## Supplemental Information

**Supplementary Figure 1:**

The presence of the tTR repressor affected tet-dependent Venus expression *in vitro*. Transductionof dissociated hippocampal cultures at DIV 7 with the standard driver AAV [1] containing rtTA and tTR (a) produced a different doxycycline dose-response curve than a driver AAV containing only rtTA and luciferase (b). Cells were co-transduced with a responder AAV with a tet-dependent Venus construct [2].

**Supplementary Figure 2:**

Comparison of different administration routes of doxycycline *in vivo*. After injection of the standard driver and the tet-Venus responder viruses [AAVs 1,2], doxycycline was administered intraperitoneally (IP) or by direct intracerebroventricular injection (ICV), either ipsi- or contra lateral to the imaging site. Shown is one example of each doxycycline administration route visualized by two-photon *in vivo* imaging. In each animal, the same cortical region was imaged over time, from prior to doxycycline administration up to 72 h thereafter (scale bar: 50 μm).

**Supplementary Figure 3:**

Monitoring and manual adjustment of breathing rates. (a) Sample recordings of GCaMP6f single cell calcium transients (top) and the corresponding breathing rate (bottom) of a mildly anesthetized mouse. Increased spontaneous calcium transients were detected at higher breathing rates/lower anesthesia. (b) Acute, manual adjustment of the isoflurane levels produced stable breathing rates across longitudinal experiments.

**Supplementary Figure 4:**

Registration of calcium movies from longitudinal imaging experiments. (a) Scheme of the registration process. The individual imaging planes of the raw movies were all registered to the initial time point (left) and the result was visualized by a maximum intensity projection across all time points (right). (b) A custom Matlab toolbox provides six different registration results of affine transformation matrices from which the user has to choose the preferred registration manually. Here, matrix I and V show good registration between the two pseudo-colored (green and purple) MIPs from two different time points. (c) MIP of registered imaging planes from eleven different time points produced a near complete overlapp.

**Supplementary Figure 5:**

Workflow for movie processing. (a) Scheme for removal of neuropil signal that typically ‘contaminates’ extraction of calcium transients. (b) After subtraction of the neuropil signal from the ROI signal, the corrected trace is an adequate representation of the somatic calcium transients. (c) Shown is a concatenation of calcium traces (movies) of the same cell at different time points. The uncorrected shifting baselines would impair spike estimation. Correction was achieved with a customized Matlab 2016b intrinsic Curve Fitting Toolbox 3.5.4.

**Supplementary Figure 6:**

Kir2.1-dependent changes of activity in individual neurons in vivo. (a) The absolute number of spikes was quantified in the same Cre+ neurons before (predox) and after (postdox) induction of Kir2.1. The number of spikes decreased for most neurons. (b) Same neurons as in (a) but showing the relative change to before for each individual cell. The red line indicates the overall average decrease (error bars: SD). (c) Control: The number of neurons with increasing or decreasing number of spikes was about the same in Cre- neurons. (d) Same neurons as in (c). The red line is close to zero suggestive of stable spontaneous activities across the imaged neuronal population.

**Supplementary Figure 7:**

Cellular specificity of transgene expression *in vivo*. (a) Injection of a Synapsin-GCaMP6f AAV into double transgenic PVcre : floxed tdTomato mice transduced a spatially restricted bolus of neurons in the S1/M1 cortex (left panel). Up to 20% of the tdTomato-positive parvalbumin neurons (center) also expressed viral GCaMP6f through the synapsin promoter (right panel; arrows, inset) as seen in the confocal image of a cortical brain slice (vertical brain surface to the right; scalebar: 200 μm).

**Supplementary Figure 8:**

Cell type-specific silencing of inhibitory parvalbumin neurons. (a) In one animal, induction of Kir2.1 in PV neurons induced a roughly ten-fold increase in spontaneous activity in the surrounding non-PV cells over the course of the experiment (error bars: SEM). Following a rapid increase in activity of the surrounding neurons, their activity decreased temporarily (hours 30 to 55) and then rebounded again to levels before the decrease (hour 25 vs. hour 80). (b) Average of the three animals in which the intermittent decrease was observed (error bars: SEM).

## Supplemental Methods

### Post-image processing and spike extraction

#### Motion correction and movie registration

Non-rigid correction for respiratory or mechanical motion drifts between individual imaging sessions was accomplished using the ImageJ Fiji distribution (Schindelin et al., 2012) and the plugin “Moco” (Dubbs et al., 2016). For the analysis from consecutive time points, the respective imaging planes had to be registered onto each other. A time point before doxycycline induction was chosen as the template image to which all the other time points were registered. To achieve a high fidelity registration of calcium imaging movies, we created a ‘Calcium Movie Registration (CaMoReg) toolbox’ based on Matlab 2016b (Mathworks). CaMoReg was developed by using the Matlab-App called “Feature-Based Image Registration 1.0.0.1 by Brett Shoelson”. CaMoReg works semi-automatically as it requires the user to choose the best registration among six different non-rigid image registration algorithm results. This App utilizes all of Matlab intrinsic Computer Vision System Toolbox parameters on an interactive user interface and displays the immediate results.

#### Calcium trace extraction

For the extraction of calcium traces from regions that were typically densely packed with GCaMP6f positive neurons, the contaminant neuropil signal was removed. For this, cell somata were manually selected in ImageJ using the maximum intensity projections of all registered time points. This ROI set was loaded for every experiment into a custom-written Matlab based algorithm that processes each single ROI separately. Each ROI was expanded by 4 pixels around the initial sROI, but not to overlap with neighboring ROIs. This created a donut-shaped, second neuropil-ROI around the initial soma-ROI. The neuropil-ROI represents ‘contaminant’ signal from dendrites and axons in the vicinity of the soma-ROI. For each frame, the average fluorescence value of the neuropil-ROI was subtracted from the average fluorescence value of the soma-ROI. After processing every ROI for every time point of the experiment, the results of the neuropil corrected calcium traces were saved for each ROI in a matrix for every imaging frame.

Fluctuating or shifting baselines of calcium signals were corrected using a Matlab 2016b script based on the intrinsic Curve Fitting Toolbox 3.5.4 with either one of four constrained curve fitting models, from rigid to flexible. The grade of flexibility was determined by the degree of the polynomial fit that was utilized for every model. A user interface was added to the script to display the result of every curve fitting model and allow the user to choose the best fit manually. The whole process was reiteratively performed on all single cells calcium traces to achieve to best possible baselines for each calcium trace.

#### Spike estimation

With the established image processing pipeline, calcium traces could be extracted from single cells and analyzed over multiple time points. To analyze the activity of neuronal cell populations and inferring their spiking patterns, the MLspike algorithm was used(Deneux et al., 2016). The algorithm was first calibrated with a published *in vivo* dataset (Chen et al., 2013) that was very similar to our recordings. In total, 16 different parameter combinations were tested to approximate spiking from calcium traces. The combination of the MLspike intrinsic parameters Hill 2.80, C(0) 0.65, and Drift 3.00 produced only a minor spike average underestimation of −1.85 % compared to patch clamp determined spiking numbers.

#### Distance and activity correlations

The lengths of all distances between neurons that only expressed GCaMP6f and those neurons that were genetically silenced (Cre+, tdTomato positive) were summed up and divided by the number of Cre+ cells. This value indicated the relative average distance each GCaMP6 positive cell had to the silenced cells that surrounded it. The calculations were performed using the Matlab extension ReadImageJROI (Muir and Kampa, 2014) to extract the positions of the respective ROIs which were determined using Fiji. The spontaneous activity before and after induction of Kir2.1 was averaged, and a ratio calculated. To avoid divisions by zero, the values were increased by 1, and all ratios above 1 were plotted as 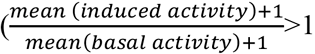. The relative average distance was correlated to the ratio of activity changes before and after induction (80 h) for all animals.

**Statistical analyses** including tests for normality were done using Prism software. Numbers are given as mean ± SD or ± SEM as indicated.

## References

Armbruster, B.N., Li, X., Pausch, M.H., Herlitze, S., and Roth, B.L. (2007). Evolving the lock to fit the key to create a family of G protein-coupled receptors potently activated by an inert ligand. Proceedings of the National Academy of Sciences of the United States of America 104, 5163–5168.

Banker, G., and Goslin, K. (1998). Culturing nerve cells, 2nd edn (Cambridge, Mass.: MIT Press).

Beard, C., Hochedlinger, K., Plath, K., Wutz, A., and Jaenisch, R. (2006). Efficient method to generate single-copy transgenic mice by site-specific integration in embryonic stem cells. Genesis 44, 23–28.

Burrone, J., O’Byrne, M., and Murthy, V.N. (2002). Multiple forms of synaptic plasticity triggered by selective suppression of activity in individual neurons. Nature 420, 414–418.

Cambridge, S.B., Geissler, D., Calegari, F., Anastassiadis, K., Hasan, M.T., Stewart, A.F., Huttner, W.B., Hagen, V., and Bonhoeffer, T. (2009). Doxycycline-dependent photoactivated gene expression in eukaryotic systems. Nature methods 6, 527–531.

Chen, T.W., Wardill, T.J., Sun, Y., Pulver, S.R., Renninger, S.L., Baohan, A., Schreiter, E.R., Kerr, R.A., Orger, M.B., Jayaraman, V., et al. (2013). Ultrasensitive fluorescent proteins for imaging neuronal activity. Nature 499, 295–300.

Deneux, T., Kaszas, A., Szalay, G., Katona, G., Lakner, T., Grinvald, A., Rozsa, B., and Vanzetta, I. (2016). Accurate spike estimation from noisy calcium signals for ultrafast three-dimensional imaging of large neuronal populations *in vivo*. Nature communications 7, 12190.

Deuschle, U., Meyer, W.K., and Thiesen, H.J. (1995). Tetracycline-reversible silencing of eukaryotic promoters. Molecular and cellular biology 15, 1907–1914.

Dogbevia, G.K., Marticorena-Alvarez, R., Bausen, M., Sprengel, R., and Hasan, M.T. (2015). Inducible and combinatorial gene manipulation in mouse brain. Frontiers in cellular neuroscience 9, 142.

Dubbs, A., Guevara, J., and Yuste, R. (2016). moco: Fast Motion Correction for Calcium Imaging. Front Neuroinform 10, 6.

Ellwardt, E., Pramanik, G., Luchtman, D., Novkovic, T., Jubal, E.R., Vogt, J., Arnoux, I., Vogelaar, C.F., Mandal, S., Schmalz, M., et al. (2018). Maladaptive cortical hyperactivity upon recovery from experimental autoimmune encephalomyelitis. Nature neuroscience 21, 1–12.

Emiliani, V., Cohen, A.E., Deisseroth, K., and Hausser, M. (2015). All-Optical Interrogation of Neural Circuits. The Journal of neuroscience : the official journal of the Society for Neuroscience 35, 13917–13926.

Feil, R., Brocard, J., Mascrez, B., LeMeur, M., Metzger, D., and Chambon, P. (1996). Ligand-activated site-specific recombination in mice. Proceedings of the National Academy of Sciences of the United States of America 93, 10887–10890.

Fossat, N., Chatelain, G., Brun, G., and Lamonerie, T. (2006). Temporal and spatial delineation of mouse Otx2 functions by conditional self-knockout. EMBO reports 7, 824–830.

Freundlieb, S., Schirra-Muller, C., and Bujard, H. (1999). A tetracycline controlled activation/repression system with increased potential for gene transfer into mammalian cells. The journal of gene medicine 1, 4–12.

Gomez, J.L., Bonaventura, J., Lesniak, W., Mathews, W.B., Sysa-Shah, P., Rodriguez, L.A., Ellis, R.J., Richie, C.T., Harvey, B.K., Dannals, R.F., et al. (2017). Chemogenetics revealed: DREADD occupancy and activation via converted clozapine. Science 357, 503–507.

Gossen, M., and Bujard, H. (1992). Tight control of gene expression in mammalian cells by tetracycline-responsive promoters. Proceedings of the National Academy of Sciences of the United States of America 89, 5547–5551.

Gossen, M., Freundlieb, S., Bender, G., Muller, G., Hillen, W., and Bujard, H. (1995). Transcriptional activation by tetracyclines in mammalian cells. Science 268, 1766–1769.

Guettier, J.M., Gautam, D., Scarselli, M., Ruiz de Azua, I., Li, J.H., Rosemond, E., Ma, X., Gonzalez, F.J., Armbruster, B.N., Lu, H., et al. (2009). A chemical-genetic approach to study G protein regulation of beta cell function *in vivo*. Proceedings of the National Academy of Sciences of the United States of America 106, 19197–19202.

Halasy, K., Buhl, E.H., Lorinczi, Z., Tamas, G., and Somogyi, P. (1996). Synaptic target selectivity and input of GABAergic basket and bistratified interneurons in the CA1 area of the rat hippocampus. Hippocampus 6, 306–329.

Hasan, M.T., Schonig, K., Berger, S., Graewe, W., and Bujard, H. (2001). Long-term, noninvasive imaging of regulated gene expression in living mice. Genesis 29, 116–122.

Hayashi, S., and McMahon, A.P. (2002). Efficient recombination in diverse tissues by a tamoxifen-inducible form of Cre: a tool for temporally regulated gene activation/inactivation in the mouse. Developmental biology 244, 305–318.

Hippenmeyer, S., Vrieseling, E., Sigrist, M., Portmann, T., Laengle, C., Ladle, D.R., and Arber, S. (2005). A developmental switch in the response of DRG neurons to ETS transcription factor signaling. PLoS Biol 3, e159.

Holtmaat, A., Bonhoeffer, T., Chow, D.K., Chuckowree, J., De Paola, V., Hofer, S.B., Hubener, M., Keck, T., Knott, G., Lee, W.C., et al. (2009). Long-term, high-resolution imaging in the mouse neocortex through a chronic cranial window. Nat Protoc 4, 1128–1144.

Howe, M.W., Feig, S.L., Osting, S.M., and Haberly, L.B. (2008). Cellular and subcellular localization of Kir2.1 subunits in neurons and glia in piriform cortex with implications for K+ spatial buffering. The Journal of comparative neurology 506, 877–893.

Kistner, A., Gossen, M., Zimmermann, F., Jerecic, J., Ullmer, C., Lubbert, H., and Bujard, H. (1996). Doxycycline-mediated quantitative and tissue-specific control of gene expression in transgenic mice. Proceedings of the National Academy of Sciences of the United States of America 93, 10933–10938.

Langford, D.J., Bailey, A.L., Chanda, M.L., Clarke, S.E., Drummond, T.E., Echols, S., Glick, S., Ingrao, J., Klassen-Ross, T., Lacroix-Fralish, M.L., et al. (2010). Coding of facial expressions of pain in the laboratory mouse. Nature methods 7, 447–449.

Lin, C.W., Sim, S., Ainsworth, A., Okada, M., Kelsch, W., and Lois, C. (2010). Genetically increased cell-intrinsic excitability enhances neuronal integration into adult brain circuits. Neuron 65, 32–39.

MacLaren, D.A., Browne, R.W., Shaw, J.K., Krishnan Radhakrishnan, S., Khare, P., Espana, R.A., and ClarkS.D. (2016). Clozapine N-Oxide Administration Produces Behavioral Effects in Long-Evans Rats: Implications for Designing DREADD Experiments. eNeuro 3.

Madisen, L., Zwingman, T.A., Sunkin, S.M., Oh, S.W., Zariwala, H.A., Gu, H., Ng, L.L., Palmiter, R.D., Hawrylycz, M.J., Jones, A.R., et al. (2010). A robust and high-throughput Cre reporting and characterization system for the whole mouse brain. Nature neuroscience 13, 133–140.

Mansuy, I.M., and Bujard, H. (2000). Tetracycline-regulated gene expression in the brain. Current opinion in neurobiology 10, 593–596.

Muir, D.R., and Kampa, B.M. (2014). FocusStack and StimServer: a new open source MATLAB toolchain for visual stimulation and analysis of two-photon calcium neuronal imaging data. Front Neuroinform 8, 85.

Pluck, A. (1996). Conditional mutagenesis in mice: the Cre/loxP recombination system. International journal of experimental pathology 77, 269–278.

Pozo, K., and Goda, Y. (2010). Unraveling mechanisms of homeostatic synaptic plasticity. Neuron 66, 337–351.

Roome, C.J., and Kuhn, B. (2014). Chronic cranial window with access port for repeated cellular manipulations, drug application, and electrophysiology. Frontiers in cellular neuroscience 8, 379.

Sando, R., Bushong, E., Zhu, Y., Huang, M., Considine, C., Phan, S., Ju, S., Uytiepo, M., Ellisman, M., and Maximov, A. (2017). Assembly of Excitatory Synapses in the Absence of Glutamatergic Neurotransmission. Neuron 94, 312–321 e313.

Saunders, A., Macosko, E.Z., Wysoker, A., Goldman, M., Krienen, F.M., de Rivera, H., Bien, E., Baum, M., Bortolin, L., Wang, S., et al. (2018). Molecular Diversity and Specializations among the Cells of the Adult Mouse Brain. Cell 174, 1015–1030 e1016.

Schindelin, J., Arganda-Carreras, I., Frise, E., Kaynig, V., Longair, M., Pietzsch, T., Preibisch, S., Rueden, C., Saalfeld, S., Schmid, B., et al. (2012). Fiji: an open-source platform for biological-image analysis. Nature methods 9, 676–682.

Shamir, M., Bar-On, Y., Phillips, R., and Milo, R. (2016). SnapShot: Timescales in Cell Biology. Cell 164, 1302–1302 e1301.

Sigler, A., Oh, W.C., Imig, C., Altas, B., Kawabe, H., Cooper, B.H., Kwon, H.B., Rhee, J.S., and Brose, N. (2017). Formation and Maintenance of Functional Spines in the Absence of Presynaptic Glutamate Release. Neuron 94, 304–311 e304.

Sohal, V.S., Zhang, F., Yizhar, O., and Deisseroth, K. (2009). Parvalbumin neurons and gamma rhythms enhance cortical circuit performance. Nature 459, 698–702.

Tillack, K., Aboutalebi, H., and Kramer, E.R. (2015). An Efficient and Versatile System for Visualization and Genetic Modification of Dopaminergic Neurons in Transgenic Mice. PloS one 10, e0136203.

Tsien, J.Z. (2016). Cre-Lox Neurogenetics: 20 Years of Versatile Applications in Brain Research and Counting. Frontiers in genetics 7, 19.

Turrigiano, G. (2011). Too many cooks? Intrinsic and synaptic homeostatic mechanisms in cortical circuit refinement. Annual review of neuroscience 34, 89–103.

Wang, L., Wang, Z., Zhang, F., Zhu, R., Bi, J., Wu, J., Zhang, H., Wu, H., Kong, W., Yu, B., et al. (2016). Enhancing Transgene Expression from Recombinant AAV8 Vectors in Different Tissues Using Woodchuck Hepatitis Virus Post-Transcriptional Regulatory Element. Int J Med Sci 13, 286–291.

Watakabe, A., Sadakane, O., Hata, K., Ohtsuka, M., Takaji, M., and Yamamori, T. (2017). Application of viral vectors to the study of neural connectivities and neural circuits in the marmoset brain. Developmental neurobiology 77, 354–372.

Xu, H.T., Pan, F., Yang, G., and Gan, W.B. (2007). Choice of cranial window type for *in vivo* imaging affects dendritic spine turnover in the cortex. Nature neuroscience 10, 549–551.

Yarmolinsky, M., and Hoess, R. (2015). The Legacy of Nat Sternberg: The Genesis of Cre-lox Technology. Annual review of virology 2, 25–40.

Zehendner, C.M., Luhmann, H.J., and Yang, J.W. (2013). A simple and novel method to monitor breathing and heart rate in awake and urethane-anesthetized newborn rodents. PloS one 8, e62628.

Zhu, P., Aller, M.I., Baron, U., Cambridge, S., Bausen, M., Herb, J., Sawinski, J., Cetin, A., Osten, P., Nelson, M.L., et al. (2007). Silencing and un-silencing of tetracycline-controlled genes in neurons. PloS one 2, e533.

