## Supplementary figures and images for "Genetic engineering for *in vivo* optical interrogation of neuronal responses to cell type-specific silencing"

### Supplementary Figure 1

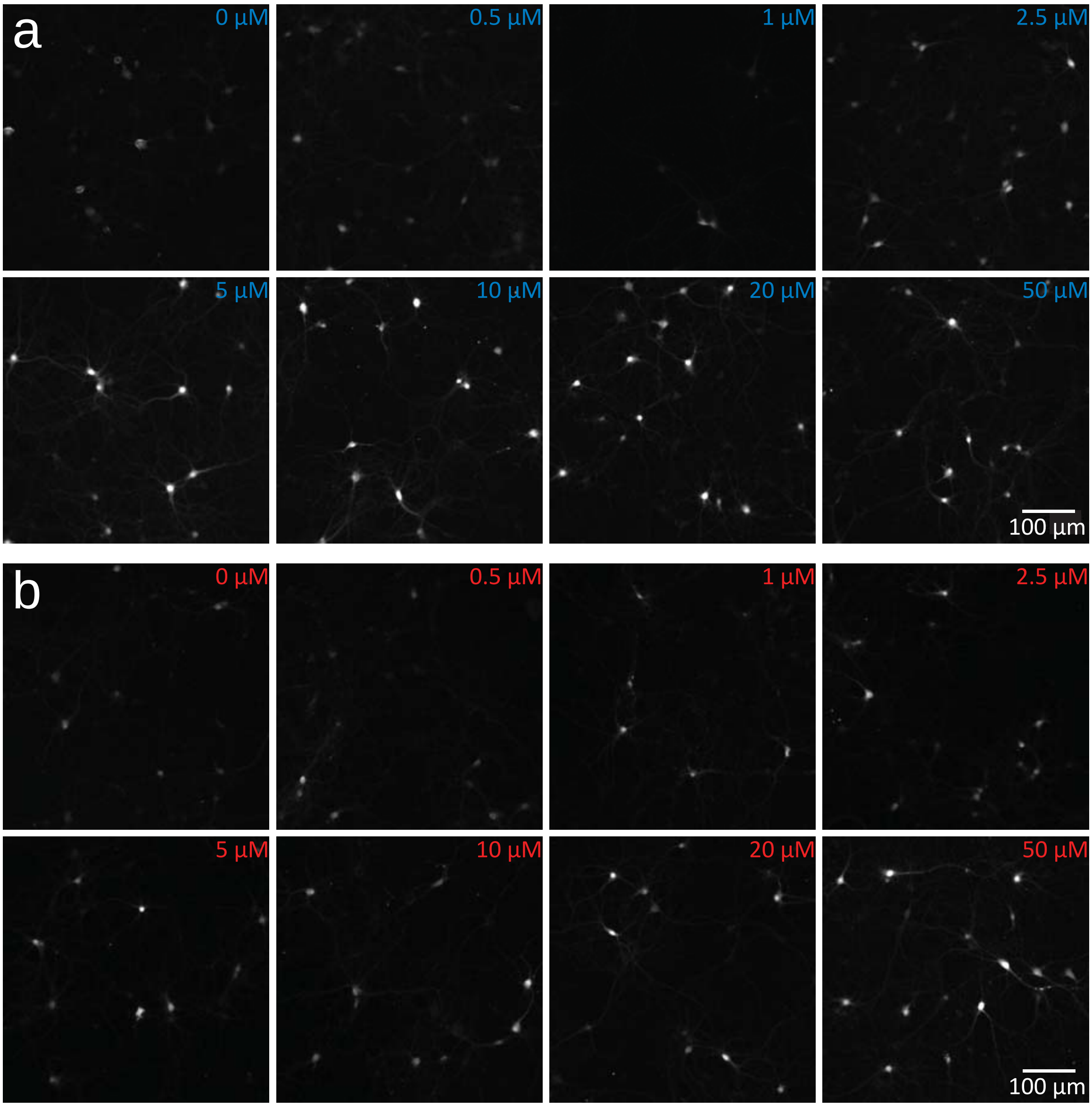


Supplementary Figure 1

### Supplementary Figure 2

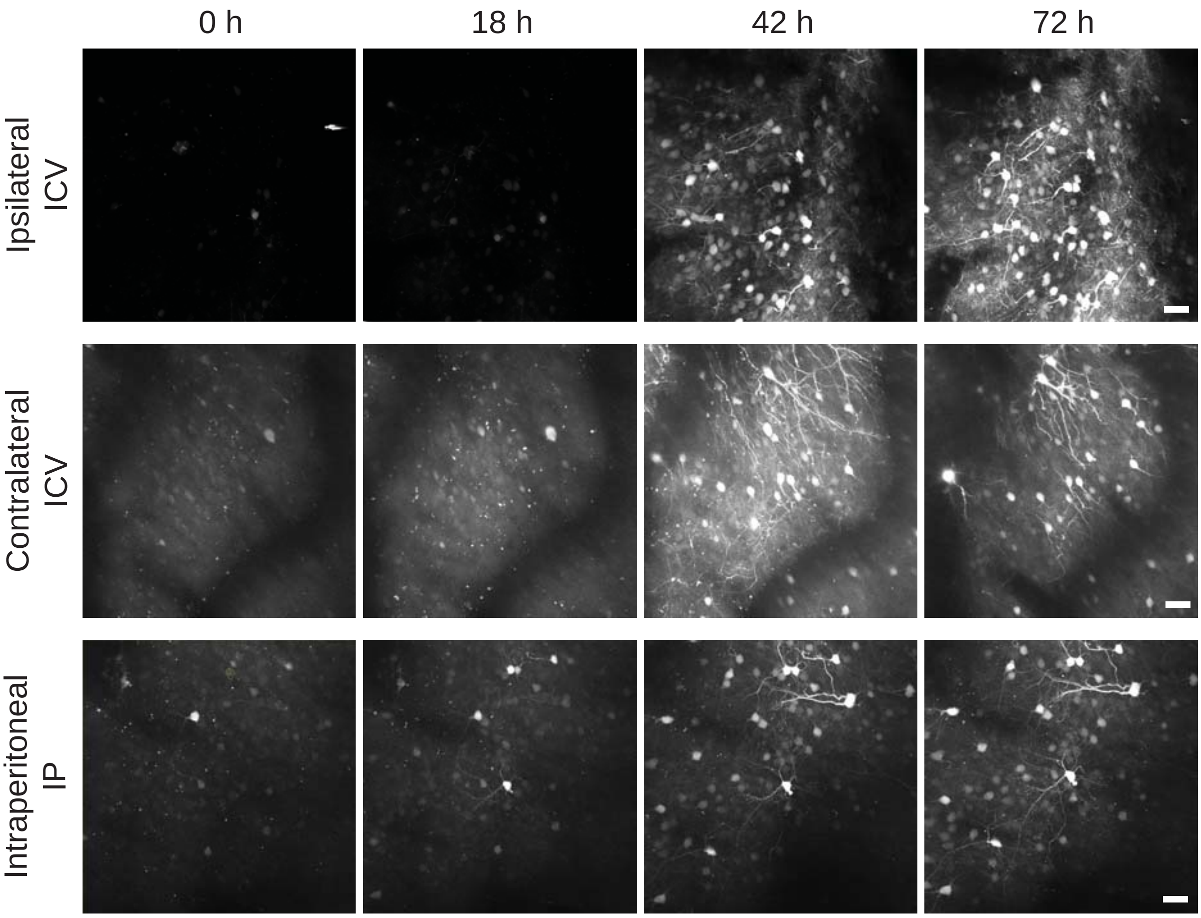


Supplementary Figure 2

### Supplementary Figure 3

a

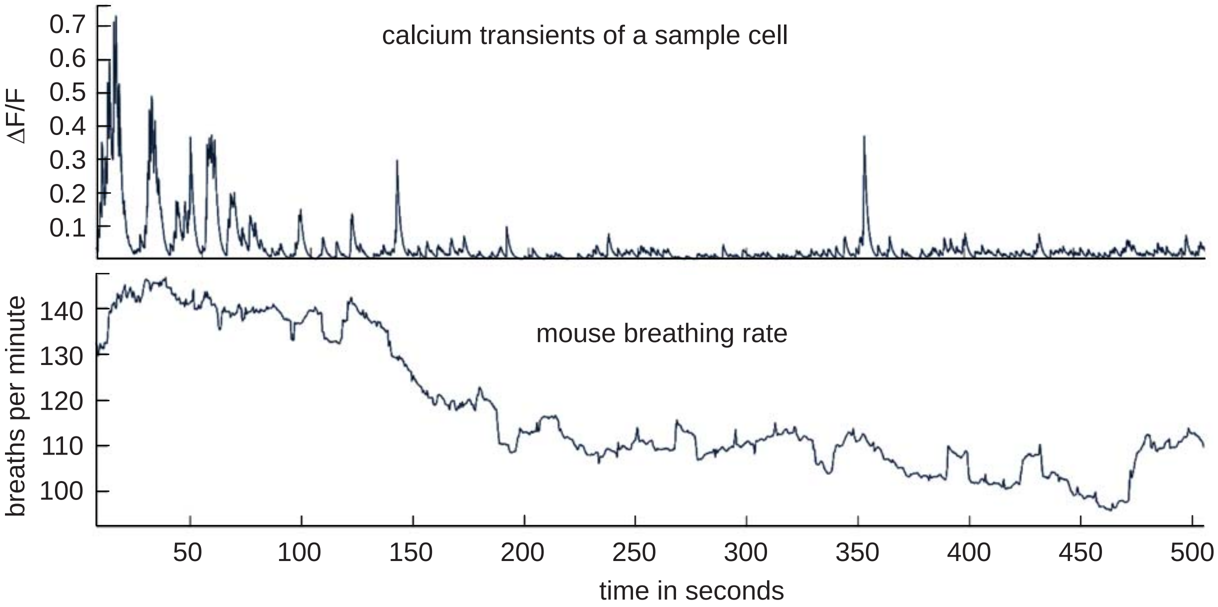


b

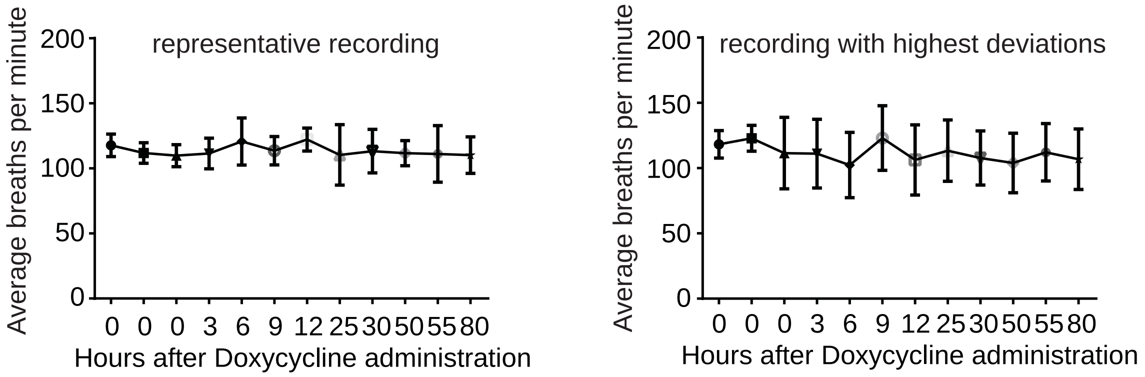


Supplementary Figure 3

### Supplementary Figure 4

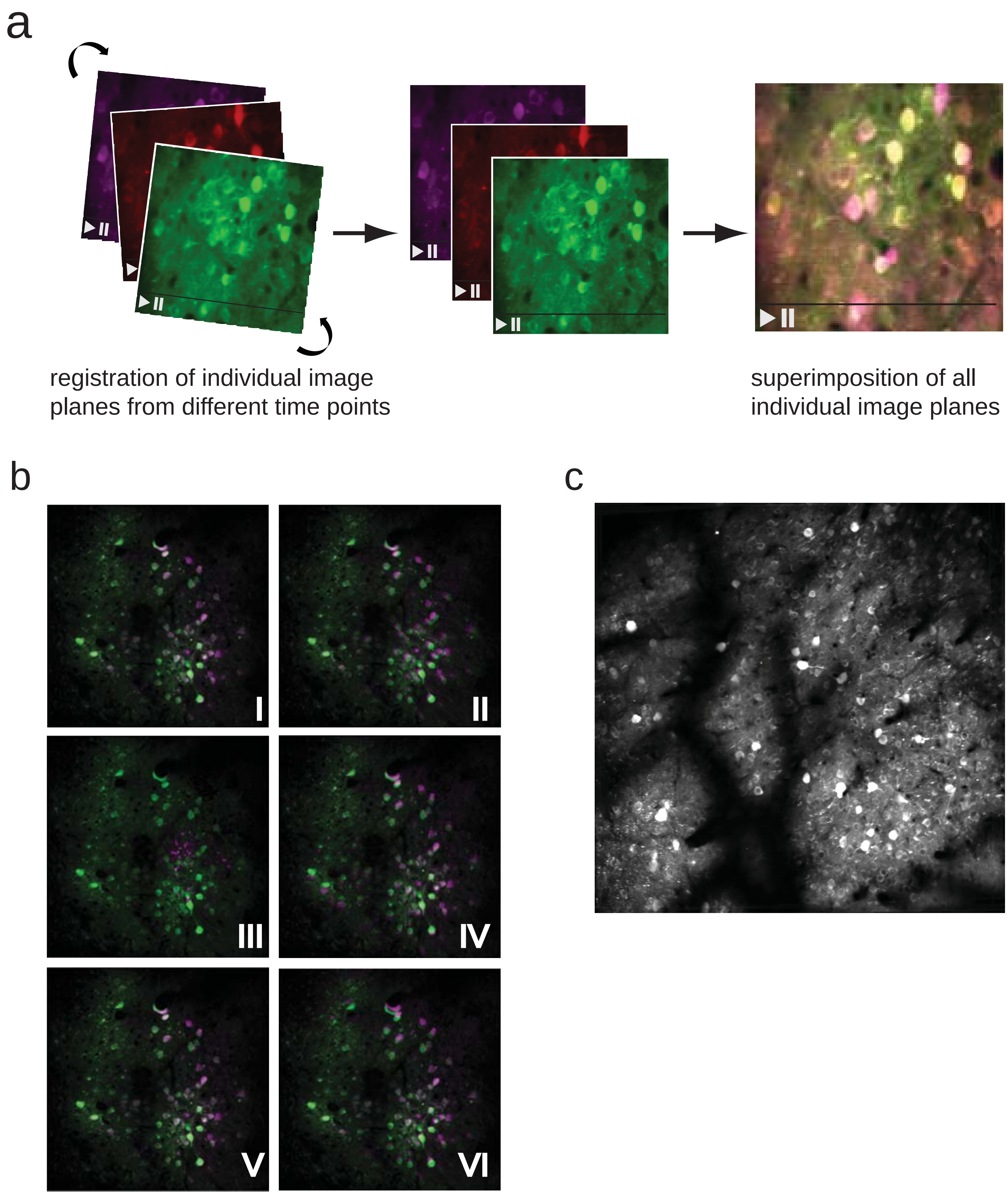


Supplementary Figure 4

### Supplementary Figure 5

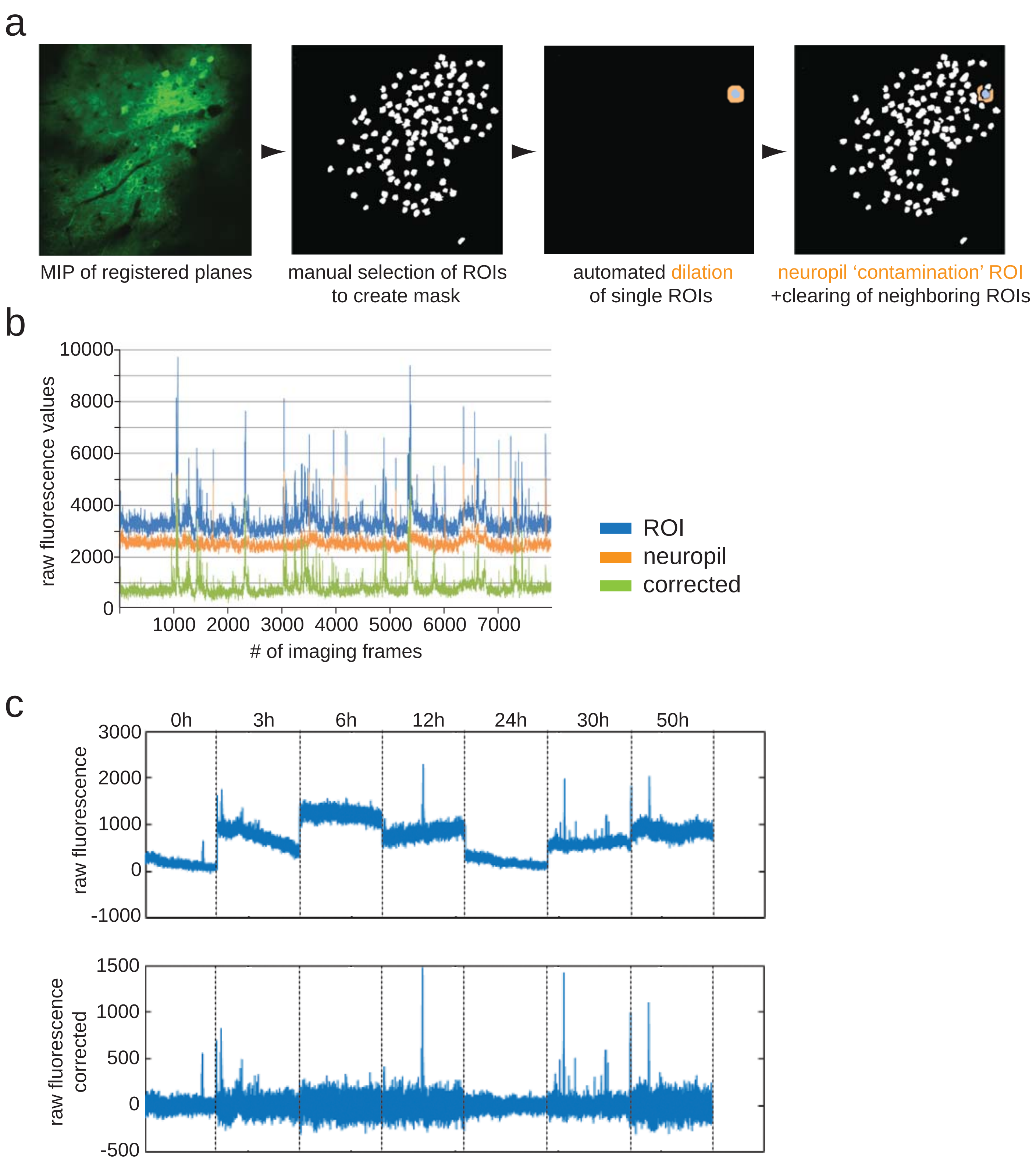


Supplementary Figure 5

### Supplementary Figure 6

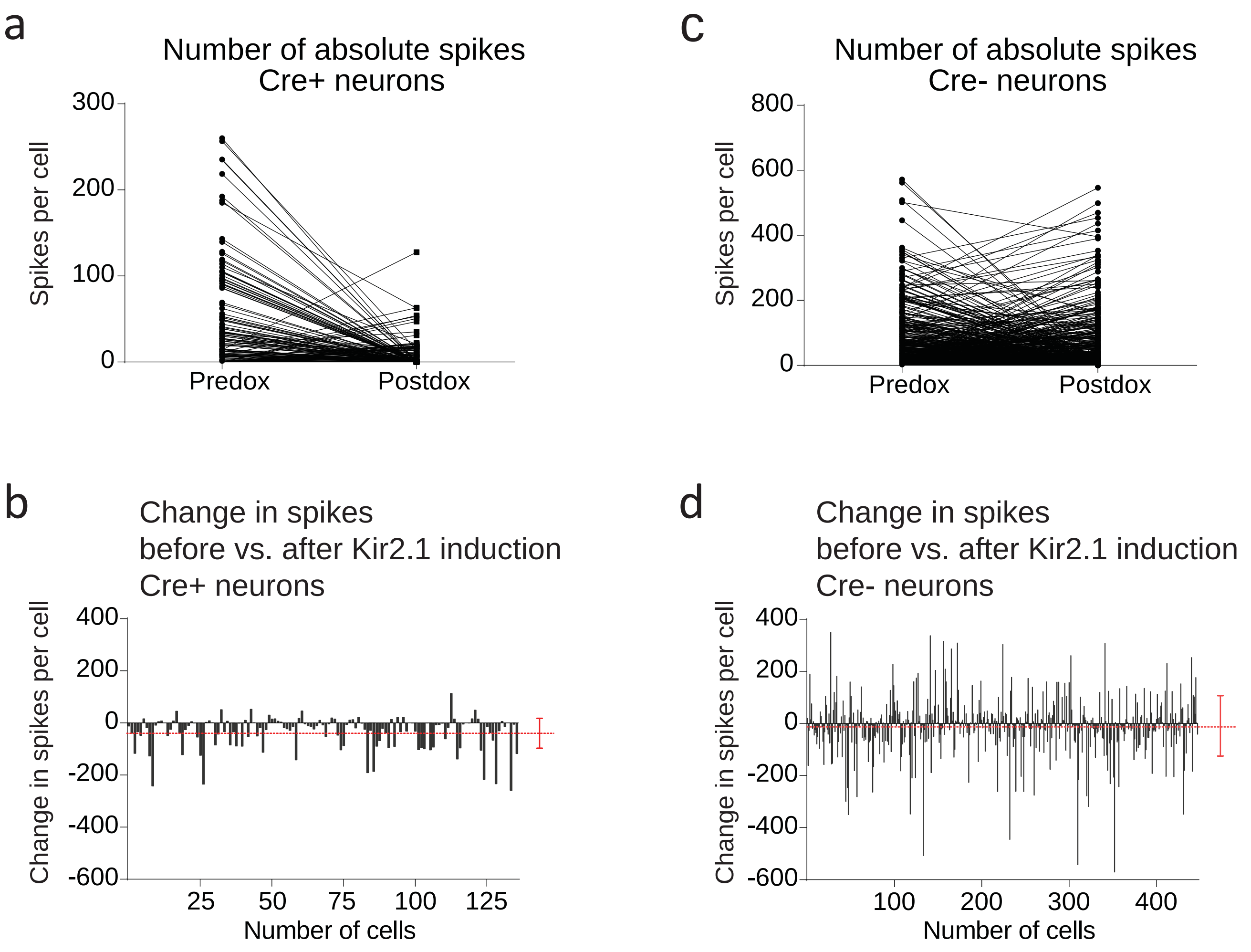


Supplementary Figure 6

### Supplementary Figure 7

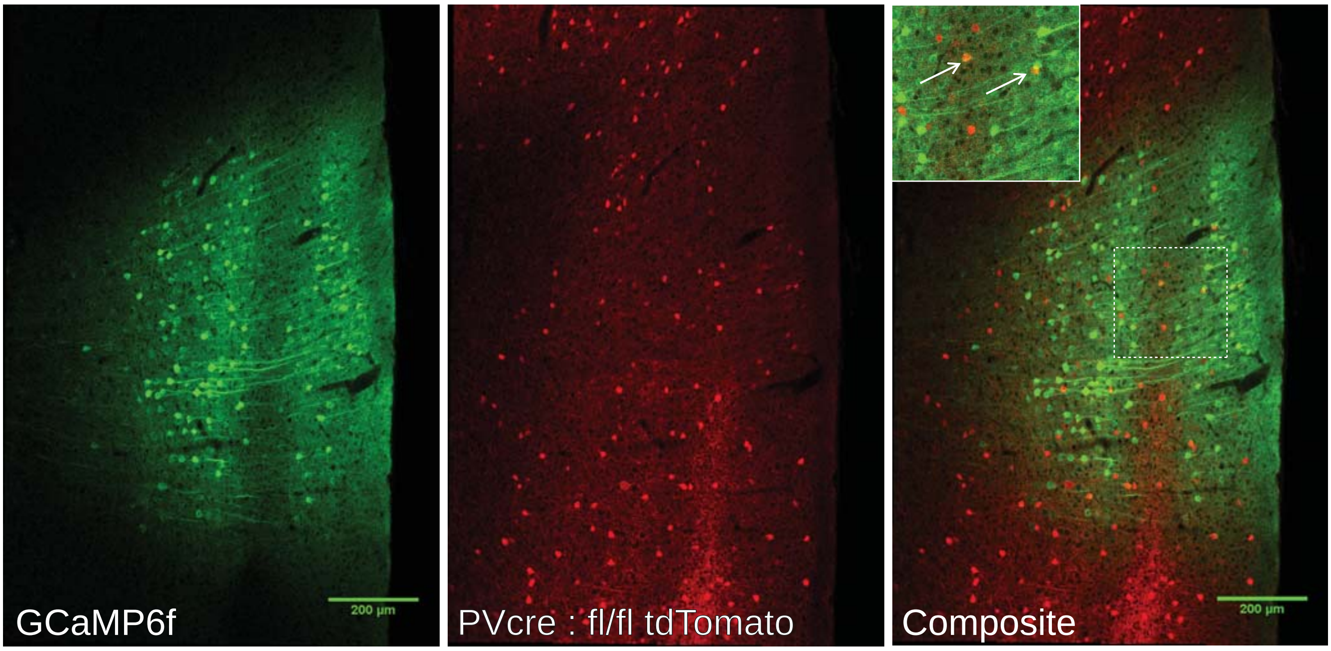


Supplementary Figure 7

### Supplementary Figure 8

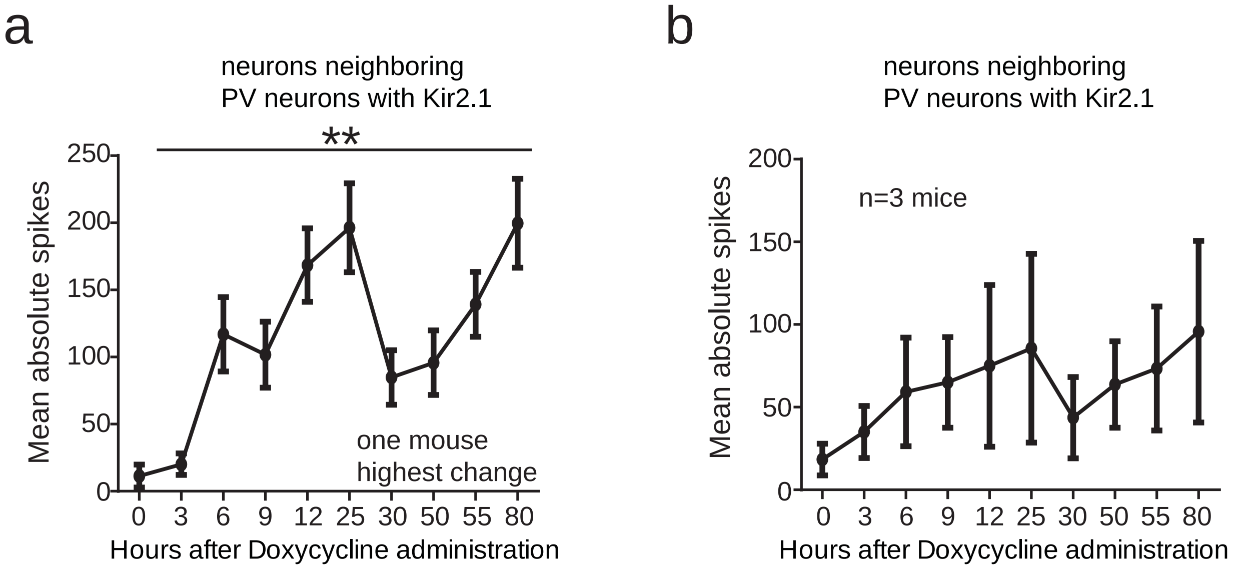


Supplementary Figure 8
